# Differential kinematic coding in sensorimotor striatum across species-typical and learned behaviors reflects a difference in control

**DOI:** 10.1101/2023.10.13.562282

**Authors:** Kiah Hardcastle, Jesse D. Marshall, Amanda Gellis, Ugne Klibaite, William Wang, Selimzhan Chalyshkan, Bence P. Ölveczky

**Author notes:** Denotes equal contribution (Kiah Hardcastle, Jesse D. Marshall).

## Abstract

The sensorimotor arm of the basal ganglia is a major part of the mammalian motor control network, yet whether it is essential for generating natural behaviors or specialized for learning and controlling motor skills is unclear. We examine this by contrasting contributions of the sensorimotor striatum (rodent dorsolateral striatum, DLS) to spontaneously expressed species-typical behaviors versus those adapted for a task. In stark contrast to earlier work implicating DLS in the control of acquired skills, bilateral lesions had no discernable effects on the expression or detailed kinematics of species-typical behaviors, such as grooming, rearing, or walking. To probe the neural correlates underlying this dissociation, we compared DLS activity across the behavioral domains. While neural activity reflected the kinematics of both learned and species-typical behaviors, the coding schemes were very different. Taken together, we did not find evidence for the basal ganglia circuit being required for species-typical behaviors; rather, our results suggest that it monitors ongoing movement and learns to alter its output to shape skilled behaviors in adaptive and task-specific ways.

## Introduction

The sensorimotor arm of the mammalian basal ganglia (BG) has long been implicated in motor control, yet which types of movements and behaviors it contributes to remains contentious^1–7^. One perspective posits that it exerts influence over a wide range of behaviors by modulating motor controllers in both brainstem and cortex^3,8–13^. Conversely, the BG circuit could function predominantly as a learning machine, specialized to support the acquisition and control of new skills^5,14–16^. Under this view, the BG would not be required for executing or specifying the details of species-typical behaviors, i.e., ones shared across members of the same species and shaped in early development^17^. These would instead be solely generated by the descending motor pathway, including dedicated pattern generator circuits in the brainstem and spinal cord^18–20^. For learned behaviors, however, the BG would act on these downstream control circuits^11,21^, affecting their dynamics in ways that improve - through learning - the kinematics of task-specific movements, making them more efficient, fluid, and aligned with the task. In this study, we directly investigate which of these hypotheses best describes BG function.

Multiple lines of evidence suggest that the BG are essential parts of the motor control machinery underlying basic movements and actions. First, they are strategically located at the nexus of the distributed motor system^22,23^, sending/receiving information to/from motor controllers in brainstem and cortex, thus allowing for continuous, if indirect, control of muscle activity^9^. Second, changes to normal BG activity, due to either disease or experimental manipulations, often results in general control deficits and can affect motor output across a range of behaviors^24–26^. Third, neural activity in the BG correlates with movement during many natural behaviors, consistent with a role in their specification and control^27–30^.

However compelling, there are plausible alternative explanations for these observations that call into question BG’s role in general motor control. For one, disease-related changes to BG activity could cause aberrant BG output that interferes with the dynamics of downstream control circuits, thus causing motor symptoms that do not accurately reflect BG function^4,31^. Consistent with this view, chronically silencing BG output can restore lost function in patients with striatal dysfunction^32–35^. Further, correlations of neural activity with a given behavior, while consistent with a control function, do not establish a causal link.

Alternatively, BG could be dispensable for basic motor control, acting primarily to shape the kinematics of task-specific learned behaviors through actions on downstream control circuits^2,4,36^ (Fig 1A-B). Evidence across species supports this perspective. In primates, for instance, disruptions to the input layer of the sensorimotor BG, the sensorimotor striatum, impair execution of learned motor sequences while leaving other movements relatively unaffected^37^. And likewise, studies in rodents likewise point to an essential role for the sensorimotor striatum (termed dorsolateral striatum, or DLS) in the acquisition and faithful execution of learned skills^38–40^. For example, in a lever-pressing task in which rats acquire stereotyped and idiosyncratic task-specific movement patterns, DLS lesions in expert animals caused a reversion to species-typical movements expressed early in learning^36,41^. Additionally, DLS activity during the task was closely linked to the animals’ movements and could account for trial-to-trial kinematic variability^36,41^, consistent with DLS specifying the details of the learned movement patterns.

**Figure 1.**
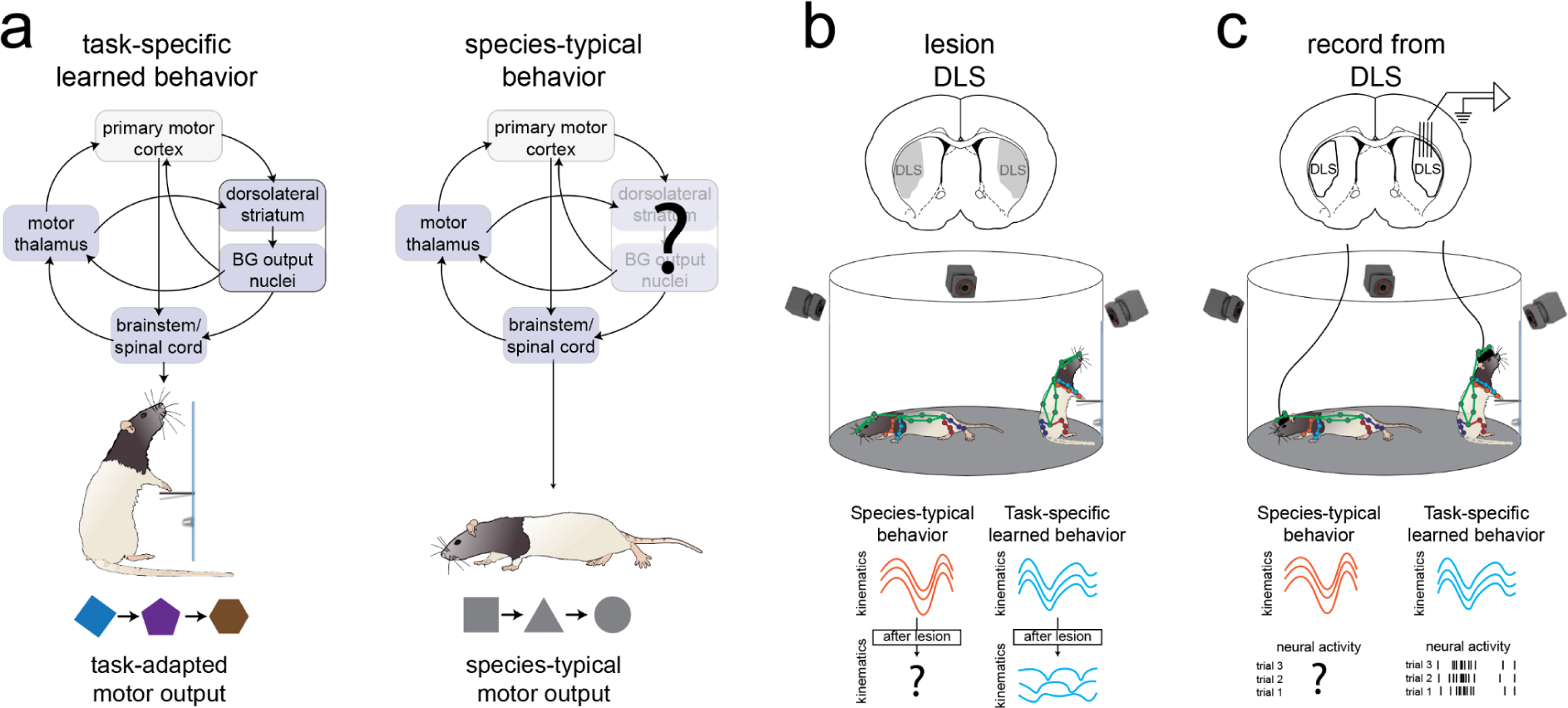
Schematic of the central hypothesis and experimental design. **a.** Schematic depicting the proposed role of the basal ganglia during species-typical behavior (right) and during execution of a learned task (left). Brain regions hypothesized to be regions essential for the behavior are in blue, whereas non-essential regions are in gray. The icons beneath the animal represent the basic movements constituting species-typical behavior (right), or adapted versions of basic movements that are specially tailored for task execution (left). Rat icons from scidraw.io, doi.org/10.5281/zenodo.3926125, doi.org/10.5281/zenodo.3926277. **b.** Schematic of lesion experiments. Top: schematic of bilateral DLS lesions. Middle: As indicated by the colored skeletons, the animal’s pose is tracked during spontaneous natural behavior and during the timed-lever pressing task. Changes to movement patterns are compared before and after DLS is lesioned. Bottom: The top row represents the animal’s movements (e.g., the height of the head) associated with a specific behavior in each domain before DLS is lesioned. Prior work has shown lesioning DLS disrupts learned movement patterns^36^; here we test the impact of lesioning DLS on species-typical movement patterns. **c.** Schematic of electrophysiology experiments. Top: Schematic of the tetrode recordings. Middle: The animal’s whole-body 3D pose, and DLS neural activity, is tracked during both exploratory and task-specific behavioral domains. While prior work has indicated that activity is locked to movement during the task (bottom right), here we explore whether DLS shows similar locking, and similar kinematic representations, during species-typical behavior (bottom left).

To distinguish BG’s contributions to the expression of learned versus species-typical behaviors, we took advantage of recent advances in behavioral tracking^42–45^, long-term neural recordings^46^, and computational ethology^47–49^ (Fig 1C). We chose rats as our subjects because they are experimentally tractable animals^50^ with well-characterized species-typical behaviors and, importantly, the species in which BG’s role in specifying and generating learned movement patterns has been most prominently established^36,38,50–53^. We view BG’s motor control function through the lens of DLS and ask whether it specifies the detailed kinematics of naturally expressed species-typical behaviors as it does for learned ones^36^.

By comparing the behavioral effects of DLS lesions on the kinematics and sequencing of natural behaviors expressed during free exploration (i.e. species-typical) versus skilled behaviors generated in a well-practiced task (i.e., learned), and probing how activity of the very same DLS neurons reflects motor output in the two domains, we achieve an “apples-to-apples” comparison of the two hypotheses elaborated above (Fig 1C). In contrast to what we have previously observed for learned movement patterns^36^, we find that DLS is dispensable for generating species-typical behaviors, including the precise kinematics of their constituent movements. However, DLS activity reflected continuous kinematics during the same exploratory behaviors that were unaffected by DLS lesions. Consistent with DLS’s differential contribution to motor output, the mapping between neural activity and whole-body kinematics was markedly different across the distinct behavioral domains. These findings suggest that while DLS activity reflects kinematics at all times, the kinematic code for learned skills is far more effective in driving downstream circuits than the one for species-typical behavior. This is consistent with BG serving a passive observer function as animals explore their environment, but gets in on the act of shaping motor output in an experience-dependent manner during tasks. Combined, these results advance a view in which BG’s primary function in motor control is to shape the detailed kinematics of skilled behaviors.

## Results

### DLS is not required for species-typical behaviors

To probe a role for DLS in general motor control, we first considered whether it is required for generating naturally-expressed movements and behaviors. To evoke these, we let rats freely explore a large (2 foot diameter) behavioral arena for 90 minutes each day. As the behaviors expressed in these sessions were similar across individuals^44^, we refer to them as ‘species-typical’^17^, contrasting them with learned behaviors acquired in a task. To capture their structure and detailed kinematics with a precision similar to task-relevant behaviors, we employed a state-of-the art 3D pose tracking system, DANNCE^43^, to track the positions of 20 anatomical keypoints in freely behaving rats (Fig S1A-C, see Methods).

To investigate DLS’s influence over species-typical behaviors and their constituent movements, we lesioned it bilaterally as in prior task-based experiments^36^. Excitotoxic lesions were performed one hemisphere at a time with a 2-week recovery between each lesion to minimize long-term effects of the procedure on downstream structures^54–56^. Histological examination confirmed substantial bilateral DLS lesions that were similar or larger than those that induced deficits in learned skill execution (Fig S2, see prior work^36^ for comparison). We recorded animals over 4 days both before and after the lesions, comparing the lesion cohort to a control group that underwent sham lesions, i.e. craniotomies without the injection of an active agent (n = 4 animals in each cohort).

To identify distinct species-typical behaviors, we used an established analysis pipeline, “MotionMapper”^48^, which we adapted for 3D keypoint data following prior work^44^. This pipeline uses dynamic postural embedding and clustering to categorize behavior with different levels of granularity^47,48^. Specifically, MotionMapper computes high-dimensional kinematic features based on continuous whole-body pose data and then embeds these features into a lower-dimensional space, creating a “behavioral map” reflecting the entire spectrum of behavior (Fig 2A, Fig S1D, see Methods). In this map, distinct locations correspond to distinct behaviors, while the density of points in that location corresponds to how often that behavior is expressed. Critically, features from all sessions and animals are embedded into the same space, allowing for straightforward comparison of behavior both within and across animals.

**Figure 2.**
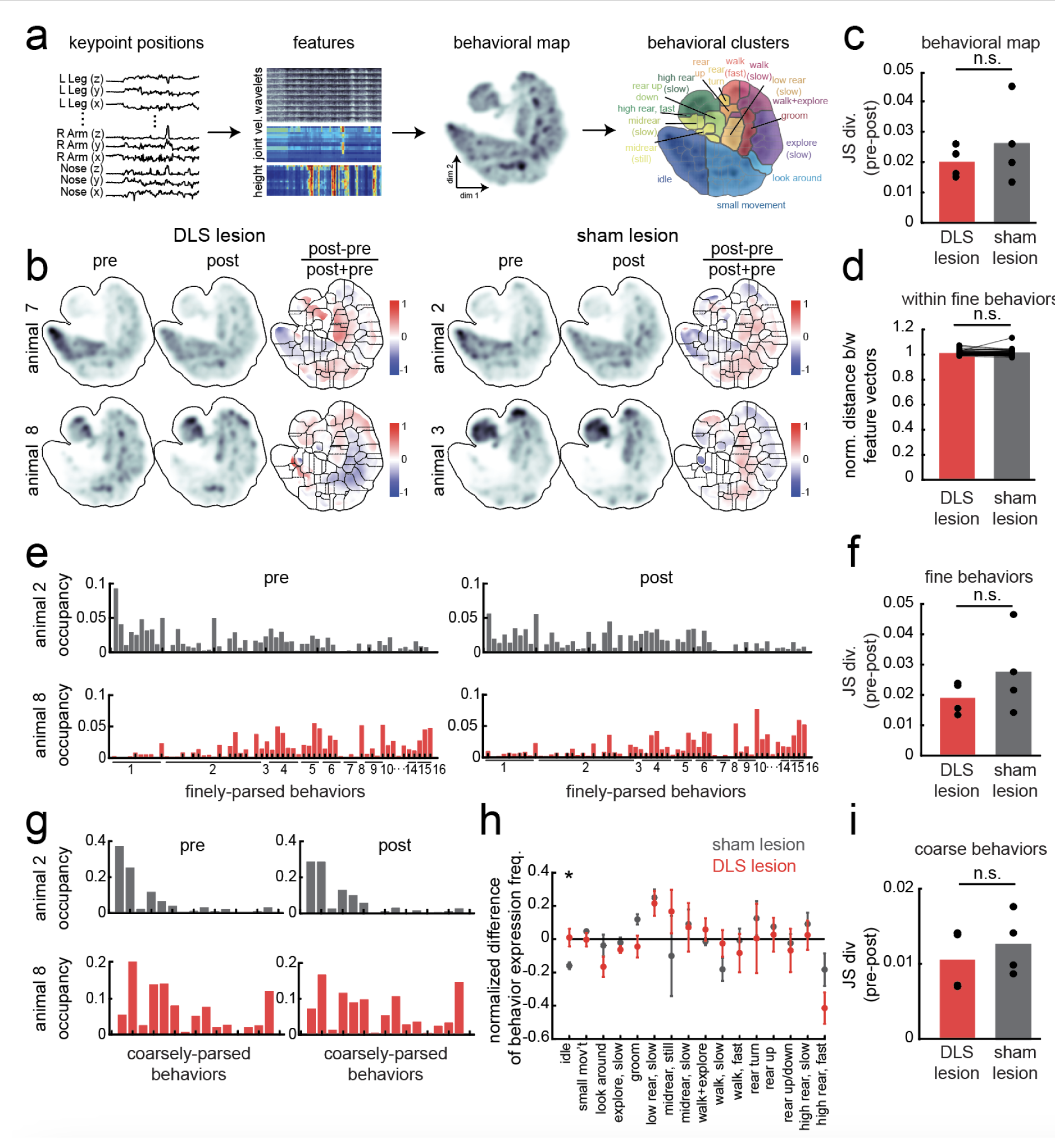
Lesioning DLS does not change the expression of species-typical behaviors. **a.** Schematic of the MotionMapper pipeline used to identify species-typical behaviors (see also Fig S1D). The labeled clusters on the right-most panel correspond to the 16 coarsely-parsed behaviors, while the gray boundaries delineate the 62 finely-parsed behaviors. **b.** The average behavioral map for two representative animals before and after DLS (left) or sham (right) lesions. Colored maps indicate the difference across maps, normalized by the sum across maps, with the fine behavioral parcellation overlaid. **c.** Comparison of the JS divergence between behavioral map distributions. There is no significant difference between cohorts (ranksum p = 0.74, n = 4 in each cohort). **d.** The average Euclidean distance between feature vectors (columns in the feature matrix in A) post-surgery compared to pre-surgery, normalized by the average Euclidean distance between feature vectors pre-surgery (see also Fig S3B-E). There was no significant difference between cohorts (averaged across n = 4 animals in each cohort, signrank p = 0.3454, n = 60, 2 fine behaviors were excluded due to low expression). **e.** Fraction of time spent expressing each finely-parsed behavior for one example rat in each cohort (corresponding to animals in B) before and after surgery. The labels on the x-axis correspond to the index of the coarsely-labeled behavior (the order in panel h). **f.** There is no difference in the JS divergence between the probability distribution of finely-parsed behavior expression across cohorts (ranksum p = 0.49, n = 4 in each group). **g.** Fraction of time spent expressing each coarsely-parsed behavior for the same example rats as in E before and after surgery. **h.** The normalized change in behavioral expression before and after surgery, quantified as (post-pre)/(post+pre). Error bars represent the SEM across animals. DLS-lesioned animals were more idle than sham-lesioned animals (n = 4, ranksum p = 0.029), although DLS-lesioned animals exhibited a smaller change compared to pre-surgery than controls. See also Fig S3G-I. **i.** There is no difference in the JS divergence between the probability distribution of coarsely-parsed behaviors between cohorts (ranksum p = 0.49, n = 4 in each group).

To parse the continuous recordings into discrete behaviors, we segmented this map to distinguish 62 “fine” behaviors using watershed clustering (Fig 2A). Under this parcellation, each cluster represented a specific variant of a species-typical behavior, such as fast versus slow walking. We further grouped these “fine” clusters into 16 ethologically relevant “coarse” behavioral types (Fig 2A).

Using these behavioral maps and categorizations, we asked whether bilateral DLS lesions induced changes in behavior. Comparing the behavioral maps before and after DLS lesions, we observed only relatively minor changes (Fig 2B, Fig S3A), especially when compared to the behavioral maps before and after the sham lesions. In fact, the differences in the behavioral densities across animals exceeded changes across lesions (mean +/-JS divergence across sessions within an animal: 0.07 +/-0.03, across animals: 0.02 +/-0.003, n = 4, ranksum p = 0.038). Moreover, the differences in the density of points across the map that we did see in DLS lesioned animals were not significantly different from sham-lesioned controls, as quantified by the JS divergence between behavioral map densities for each cohort (Fig 2C).

While this result suggests that DLS lesions influence neither the kinematics nor the frequency of species-typical behaviors, it is possible that the confound in how these aspects of behavior - movement kinematics and their frequency of expression - contribute to the map obscures a true effect (as they both contribute). To thus probe whether there are differences in movement kinematics within a given behavior, we considered whether the detailed movement patterns associated with any of the 62 “fine” behaviors were altered by DLS lesions. To do this, we examined whether the set of high-dimensional features, which correspond to the moment-by-moment kinematics of a defined behavior, changed more after lesioning DLS than after sham-lesions. Yet consistent with DLS having no essential role in shaping movement kinematics of species-typical behaviors, the features were not significantly altered by DLS lesions (Fig 2D, Fig S3B-E). DLS lesions also did not affect the distribution of embedded points within each fine behavior, i.e., the low-dimensional embedding of the feature vectors, any more than sham lesions (Fig. S3F). Lastly, we verified that the kinematics remained intact by evaluating the amplitude and velocity (vigor) of grooming and speed of locomotion, two behaviors for which these movement features can be readily quantified. Consistent with our prior results, we observed no significant lesion-induced change to the frequency or amplitude of limb movement in detected grooming bouts (Fig S4A-B) nor was there a change in the average running speed when compared to sham-lesioned animals (Fig S4C-D).

Given that DLS does not appear to contribute significantly to kinematics, we next considered whether DLS lesions may affect the degree to which species-typical behaviors are expressed, as has been previously reported in mice^28^. To probe this, we examined how frequently each of the 62 finely-parsed or 16 coarsely-parsed behaviors were expressed before and after lesions (DLS and sham). We found that for both fine and coarse behavioral parcellations, frequency of behavioral expression was relatively unchanged (Fig 2E, G), with similar deviations in distributions observed for both DLS- and sham-lesioned animals (Fig 2F, I, Fig S3G-I).

### Direct comparison of effects induced by lesioning DLS on learned and species-typical behaviors

Our results thus far indicate that DLS lesions do not affect the kinematics or frequency of species-typical behaviors. As this is in stark contrast to our prior work on highly stereotyped learned movement patterns^36,38,51^, we developed an analysis pipeline to directly, in an ‘apples-to-apples’ manner, compare the effects of DLS lesions on stereotyped species-typical behaviors and similarly precise movement patterns observed after lengthy training on a specific motor task. We utilized a previously collected dataset from rats in which their movements were carefully tracked as they performed a well-practiced learned movement pattern acquired in our timed lever-pressing task^5736^. To compare stereotyped movements across learned and species-typical behaviors, we identified “template movements” in both behavioral domains and assessed whether DLS lesions reduced their expression. Note that, as long as the behaviors are well tracked, this analysis is agnostic to which category of behavior the “template” is derived from (see Methods).

To re-characterize the effects of DLS lesions on learned movements in the timed lever-pressing task using this template-based analysis, we analyzed a dataset in which the 2D positions of both hands were tracked in 6 rats trained to asymptotic task performance^36^ (Fig 3A). Since the hand pressing the lever is largely constrained to move in 2D^36,57^, this analysis captures the essential aspects of the learned kinematics. The template was chosen as the most frequently expressed 1-second movement pattern in the task prior to lesions (Fig 3B). We quantified the frequency of this movement before and after DLS lesions (Fig 3C, see Methods). Consistent with prior analyses, DLS lesions completely halted the expression of movement trajectories similar to the pre-lesion template, even when considering trials with inter-press intervals similar to those used to derive the template (Fig 3C-D). Lesions of the neighboring dorsomedial striatum (DMS), in contrast, had no impact on the frequency with which the template behavior was expressed (Fig 3C-D), underscoring - yet again - the distinct roles of DLS and DMS in generating learned movement patterns ^37,58,59^.

**Figure 3.**
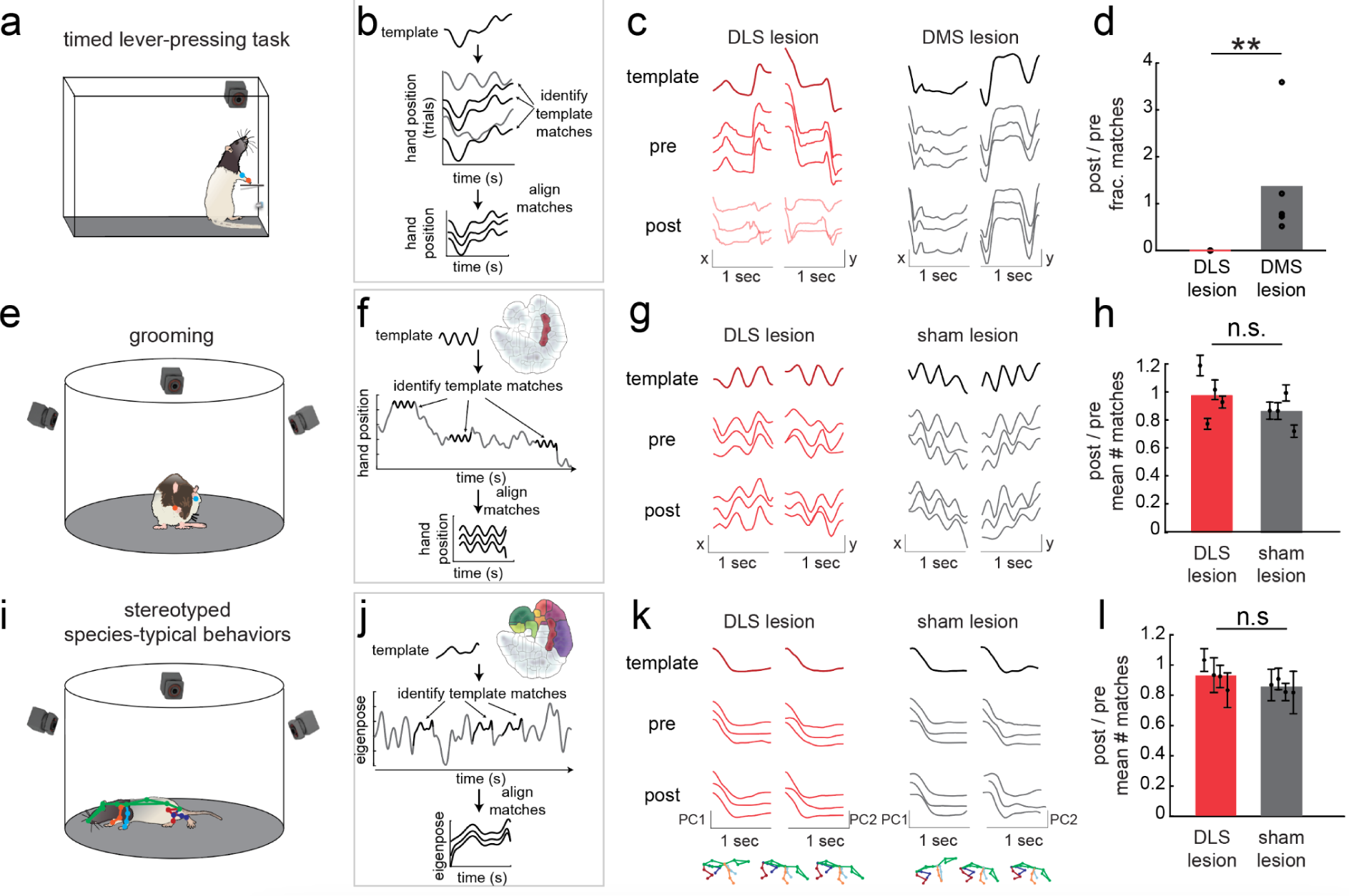
DLS lesions affect the movement kinematics of learned but not species-typical behaviors. **a.** Schematic of the timed lever-pressing task, in which rats are trained to press a lever with an inter-press-interval (IPI) of 700ms. 2D hand positions were tracked using DeeperCut^65,66^. Rat icon from scidraw.io, doi.org/10.5281/zenodo.3926125. **b.** Procedure to identify template matches across trials based on a pre-specified template. The template was identified as the most common lever-tapping movement in the trials immediately before lesion. **c.** Example templates (top) and template matches (bottom) for a single hand for two example animals before and after DLS (left) or DMS (right) lesions. Only trials in which the IPI was within the 650-750 ms range were considered. Prior to any lesion, 44% and 36% (DLS and DMS, respectively) of trials matched the template. The template matches after the DLS lesion (bottom left) did not exceed the template-matching criteria; shown instead are three of the best-matching trials. All data is from prior work^36^. **d.** The fraction of the first 5000 trials after surgery with an IPI between 650-750ms in which the animal expressed a template match compared to before surgery. DLS-lesioned animals did not exhibit any template matches after lesion, compared to DMS-lesioned animals which exhibit similar expression rates of post-lesion template matches (ranksum p = 0.004, n = 6 DLS-lesioned animals, n = 5 DMS-lesioned animals). **e.** Schematic illustrating grooming. Rat icon from scidraw.io, doi.org/10.5281/zenodo.3926223. **f.** Procedure to identify template matches across spontaneous behavior based on a pre-specified grooming template. Grooming templates were generated from behaviors expressed in the colored region of the map (see Methods). **g.** Similar to C, but for grooming movements. Top row (template): example 1-second grooming templates, shown as the x- and y-position of one hand. Middle row (pre): Three example template matches prior to surgery. Bottom row (post): Three example template matches after DLS (left) or sham (right) lesion. **h.** The ratio of template matches post-lesion compared to pre-lesion. Error bars correspond to the SEM across all templates for each animal. There was no significant difference between cohorts (no difference across groups in linear mixed-effect model with p = 0.24). Errorbars are randomly spaced along the x-axis for clarity of presentation. **i.** Schematic illustrating whole-body behaviors. Rat icon from scidraw.io, doi.org/10.5281/zenodo.3926277. **j.** Similar to F, but for whole-body behaviors. Templates were generated from the behaviors expressed during the colored region of the map, which correspond to active movements (‘walking’ but not ‘idle’, ‘small movement’, etc.). **k.** Similar to G, but showing keypoint data projected onto the top two PCs (referred to elsewhere as ‘eigenposes’, shortened here to ‘PC1’ and ‘PC2’). **l.** The ratio of template matches post-lesion compared to pre-lesion. Error bars correspond to the SEM across all templates for each animal. Error bars are randomly spaced along the x-axis for clarity of presentation. There was no significant difference between cohorts (no difference across groups in linear mixed-effect model with p = 0.08, trending towards a higher expression of template matches after surgery for DLS-lesioned animals).

For species-typical behavior, we identified templates associated with distinct and stereotyped naturalistic movements in intact rats. Initially, we focused on grooming behaviors (Fig 3E) as they are explicitly forelimb-dependent, just like the learned behavior in the timed-lever pressing task. Additionally, grooming sequences have been studied extensively in terms of striatal function, with studies suggesting a role for DLS in their sequencing^27,30,60–62^. To probe the degree to which their kinematics are affected by DLS lesions, we identified a set of pre-lesion grooming templates using the egocentrically-defined 3D positions of the left and right wrists during MotionMapper-identified grooming bouts (n = 50 ± 23 templates per animal, mean ± std, see Methods for details on template identification). Grooming templates were selected to be 1-second long (i.e., similar in length to the task-relevant movement patterns), and thus consisted of several circular movements. We quantified the number of matches per template by cross-correlating each template with the wrist positions during all sessions prior to surgery^63,64^ (Fig 3F). Note that here, unlike in the task, a wide variety of templates were considered, as rats can groom in many different ways (Fig S5A-B).

In contrast to the task-relevant template, we observed no significant change in the rate of expression of the pre-lesion grooming templates following DLS lesions compared to sham lesions (Fig 3G-H, Fig S5A-B). To further probe whether our conclusions generalized to other species-typical behaviors, we conducted a broader template-matching analysis based on 1-second long whole-body movements (Fig 3I-J, Fig S5C-D, see Methods). Unique and robustly expressed templates were identified from the 62 clusters in the behavioral map (mean ± std n = 56 ± 28 templates per animal). Similar to grooming, we observed a comparable number of matches per template before and after lesions (Fig 3K-L).

Combined, these results indicate that DLS is dispensable for the normal expression of species-typical behaviors, from their fine-grained kinematic details to their macroscopic statistical patterns of expression. This is not consistent with the DLS, or the BG more generally, being an essential cog in the motor control machinery underlying species-typical behaviors – in stark contrast to our previous results on task-specific learned motor patterns, which we further corroborated here (Fig 3A).

### DLS is not required for normal sequencing during free exploration

While DLS may not specify the detailed movements of species-typical behaviors, recent work has proposed that it may be essential for sequencing the movements and actions of naturalistic behaviors^28,61^. To examine this, we probed the transition matrices before and after surgery for both cohorts (Fig S6A-B). In contrast to prior work^28,61^, we did not find that lesioning DLS significantly impacted the transitions between behaviors relative to sham lesioned animals (Fig S6C-D). Moreover, it did not change the entropy of the behavioral transition matrix as compared to sham-lesioned animals (Fig S6E-L). This was the case whether the behaviors were parsed at the coarse or fine level, whether we used MotionMapper (Fig S6E-H) or Keypoint-Moseq to discretize behavior^67^ (Fig S6I-L), or when we only considering grooming (identified using MotionMapper, Fig S6M-N). These findings suggest that the DLS is not critical for sequencing spontaneous behaviors. Taken together, we did not find any evidence that DLS plays a significant role in determining motor output for spontaneously-expressed species-typical behaviors as recorded in freely behaving rats exploring a large arena.

### Firing statistics of DLS neurons differ across behavioral domains

If DLS indeed contributes to movement kinematics in one domain but not the other, as our lesion results suggest, this functional distinction should be reflected in the activity of its neurons. Based on prior work in which context-dependent differences in neural activity could account for differences in functional contributions^59,68–74^, we considered two main possibilities. First, differences in the statistics of neural spiking in DLS could plausibly affect its ability to influence dynamics in downstream circuits, and hence behavior. For example, if learning increases overall activity or leads to sparser and burstier firing patterns^70^, DLS’s efficiency in modulating downstream circuits and behavior could increase^70,75^. Similarly, if neurons become more strongly tuned to task-relevant movements over learning, they could be more effective in driving dynamics in downstream circuits^71,72^.

Directly probing these possibilities requires recording from the same neurons in the same animal performing both a learned skill and engaging in exploratory, i.e., species-typical, behavior. To this end, we implanted 16-tetrode arrays in DLS in rats who had reached expert performance in the timed lever-pressing task (n = 4 rats, Figure S7). To facilitate comparisons to species-typical behaviors, we also recorded in-between task sessions as animals freely explored the behavioral arena (Fig 4A). We recorded a total of 15 experimental sessions across all animals, each lasting 8-37 hours and comprising 2-3 training sessions of the timed lever-pressing task. Cells were linked across sessions using a custom spike-sorting and tracking software^46^ (n = 553 cells across all animals during the 15 sessions) and were categorized by their waveform as either spiny projection neurons (SPNs, n = 445), fast spiking interneurons (FSIs, n = 38), or other (n = 70) (see Methods).

**Figure 4.**
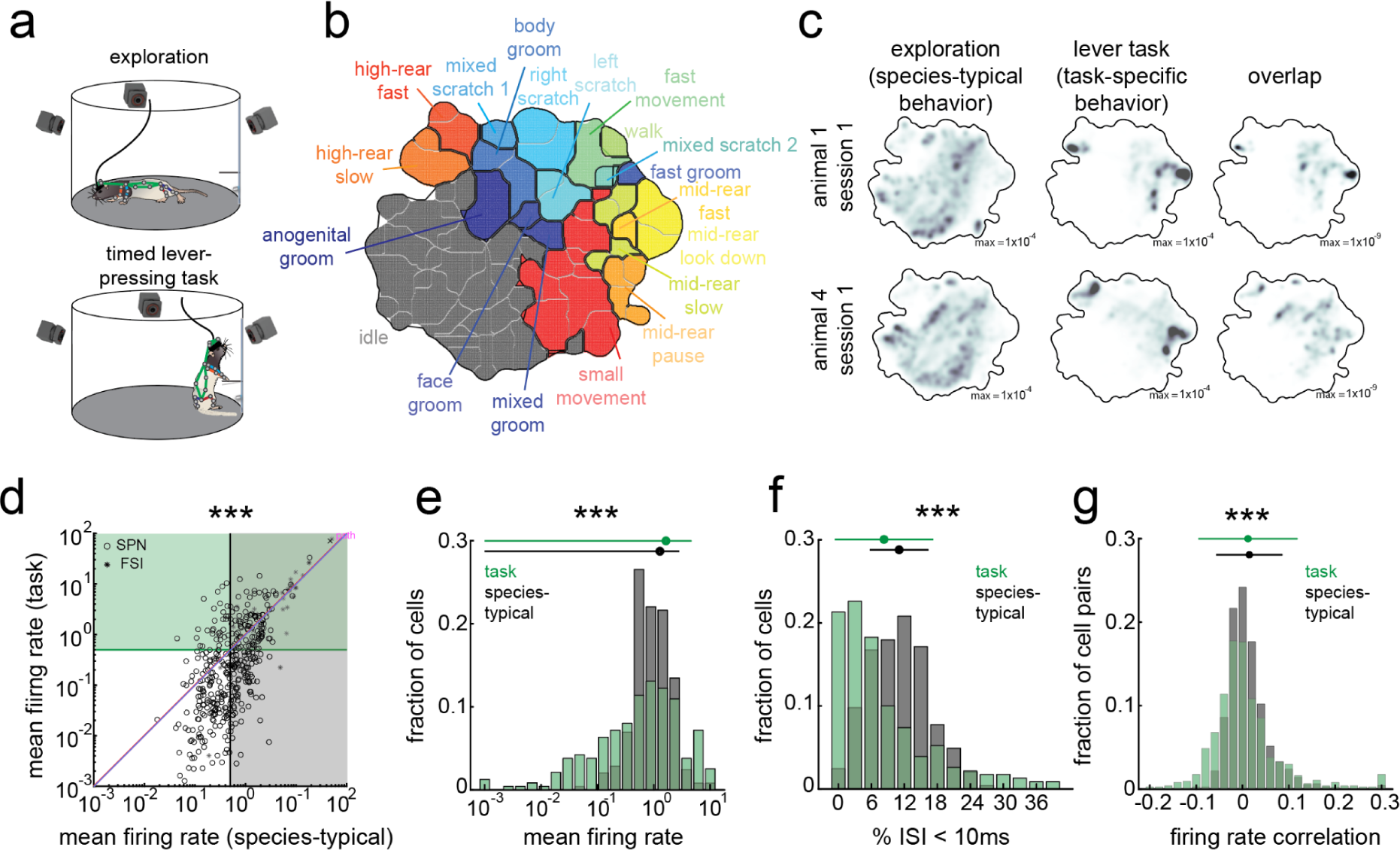
Task-specific modulation of firing rate and bursting. **a.** Depiction of the recording environment for both exploratory (top) and task-specific (bottom) behaviors. The arena was surrounded by 12 cameras to track the 3D location of 20 retroreflective markers on the animal’s body. Rat icons from scidraw.io, doi.org/10.5281/zenodo.3926125, doi.org/10.5281/zenodo.3926277. **b.** The behavioral map produced by MotionMapper for all behavioral recording sessions. Boundaries between finely-parsed behaviors (n = 74) are marked by lines, while coarsely-parsed behaviors (n = 19) are denoted by distinct colors and labels. **c.** The occupancy in the behavioral map during exploratory sessions (left) and task sessions (middle) for example recordings in two animals. The overlap between the maps is shown on the right. The numbers in the bottom-right of each panel indicate the fraction of time spent in that pixel for the darkest color. **d.** Average activity during the task-relevant or exploratory behavior for SPNs (circles) and FSIs (stars), computed across coarse behaviors with sufficient occupancy (see Methods). SPNs exhibited overall higher activity during species-typical behaviors than during the task (SPN n = 445, FSI n = 38, SPN signrank p < 0.0001, FSI signrank p = 0.46). Average activity was correlated across the two domains for both cell types (SPN: p < 0.0001, correlation coefficient = 0.72, FSI: p < 0.001, correlation coefficient = 0.97). **e.** The distribution of mean firing rates for SPNs active in either domain (in a shaded portion of panel d) during the species-typical behavior (gray) or the task (green). Plotted above the distribution are the mean +/-standard deviation of the distribution. The distribution of firing rates during the task had a significantly larger variance than during species-typical behavior (n = 245 for both groups, F-test p < 0.0001). See Fig S8A for FSIs. **f.** The distribution of bursting in SPNs, quantified as the percentage of interspike intervals less than 10 ms, for activity during the species-typical behavior (gray) or the task (green), with the mean +/-standard deviation plotted above as in d. The distribution during the task had a significantly larger variance than during species-typical behavior (n = 245 for both groups, F-test p < 0.0001). See Fig S8B for FSIs. **g.** The distribution of correlation coefficients computed between smoothed spike trains across all SPN cell pairs, with the mean +/-standard deviation plotted above. For each domain, only SNs active in that domain were considered (i.e., all SPNs in the gray zone, or green zone, of Fig 4d). The distribution of correlation coefficients during the task had a significantly larger variance than during species-typical behavior (task n = 1833, species-typical n = 742, F-test p < 0.0001). See Fig S8C for FSIs.

To accurately track the full spectrum of whole-body 3D kinematics in both behavioral domains, we employed a marker-based system (‘CAPTURE’) to register the 3D positions of 20 keypoints on the animal’s head, trunk, and limbs ^44^ (Fig 4A). We again used MotionMapper to segment the animal’s continuous movements across both domains into discrete behaviors (Fig 4B). Comparing the behavioral maps for the task-related and exploratory behaviors revealed distinct yet partially overlapping sets of kinematic motifs (Fig 4C). Thus, while the behavioral repertoire is quite different across the domains as would be expected, there is still some overlap, e.g., the animals rear in both domains. Importantly, the overlap allows us to compare the neural code for similar movements across learned and species-typical behaviors.

To probe the neurophysiological correlates for DLS’s domain-specific contributions, we first compared the average firing rate of its neurons across the domains. If firing rate is a proxy for a neuron’s engagement in a behavior, we should see less activity in DLS neurons during exploratory, i.e. species-typical, behaviors. However, this prediction was not borne out by the data. SPNs (but not FSIs) had slightly *higher* average firing rates during exploratory behavior than during the task (Fig 4D). Intriguingly, a reduction in overall activity during the expression of a motor skill aligns with observations in human and mouse motor cortex^70,72,76^, in which activity is reduced with learning. An interpretation of this result is that population-level activity diminishes with learning as neural circuits become more specialized^70,72^, leading to a smaller but more effective population of neurons driving skilled movement. To determine whether this can account for the domain specificity we see, we focused on cells that were active (average firing rate > 0.5 Hz) in at least one domain. Intriguingly, mean firing rates during the task were more widely distributed (SPNs: Fig 4E, FSIs: Fig S8A), with neurons more likely to be either very quiet or very active as compared to activity during exploratory behavior, thus consistent with this perspective.

To further probe the firing statistics of DLS neurons across domains, we examined the sparseness of neural activity, both at the level of individual neurons and across the populations. If cells concentrate their activity to specific phases of the behavior, either in terms of bursts or by correlating their firing with other neurons — it could make them more effective in influencing dynamics in downstream circuits^77^. Consistent with this, we found that cells (SPNs: Fig 4F, but not for FSIs: Fig S8B) were indeed more bursty during the task, as assessed by the percentage of interspike intervals < 10 ms. The distribution of correlation coefficients between spike trains of pairs of simultaneously recorded cells was also significantly broader (SPNs: Fig 4G, FSIs: Fig S8C). Thus, a larger subset of cells exhibited more high-frequency bursts and higher degree of correlated activity during task sessions, potentially accounting for some of the domain-specificity we see.

### DLS encodes kinematics to a greater extent during learned behavior

Since DLS contributes to kinematics only for task-relevant learned movements (Fig 2-3), we speculated that the statistical difference in firing patterns could reflect a stronger kinematic tuning to learned behaviors. To probe this, we first took a population-level perspective and asked whether fewer active neurons (denoted in Fig 4D) significantly reflected movement kinematics during exploratory, species-typical behaviors. To assess this, we employed a generalized linear modeling (GLM) framework, which allows us to quantify the number of neurons whose activity can be sufficiently predicted by whole-body pose and movement (Fig 5A). GLMs were fit to each neuron using spline functions relating to the eigenpose and eigenpose velocity, providing a highly expressive model with relatively few parameters (see Methods for more detail).

**Figure 5.**
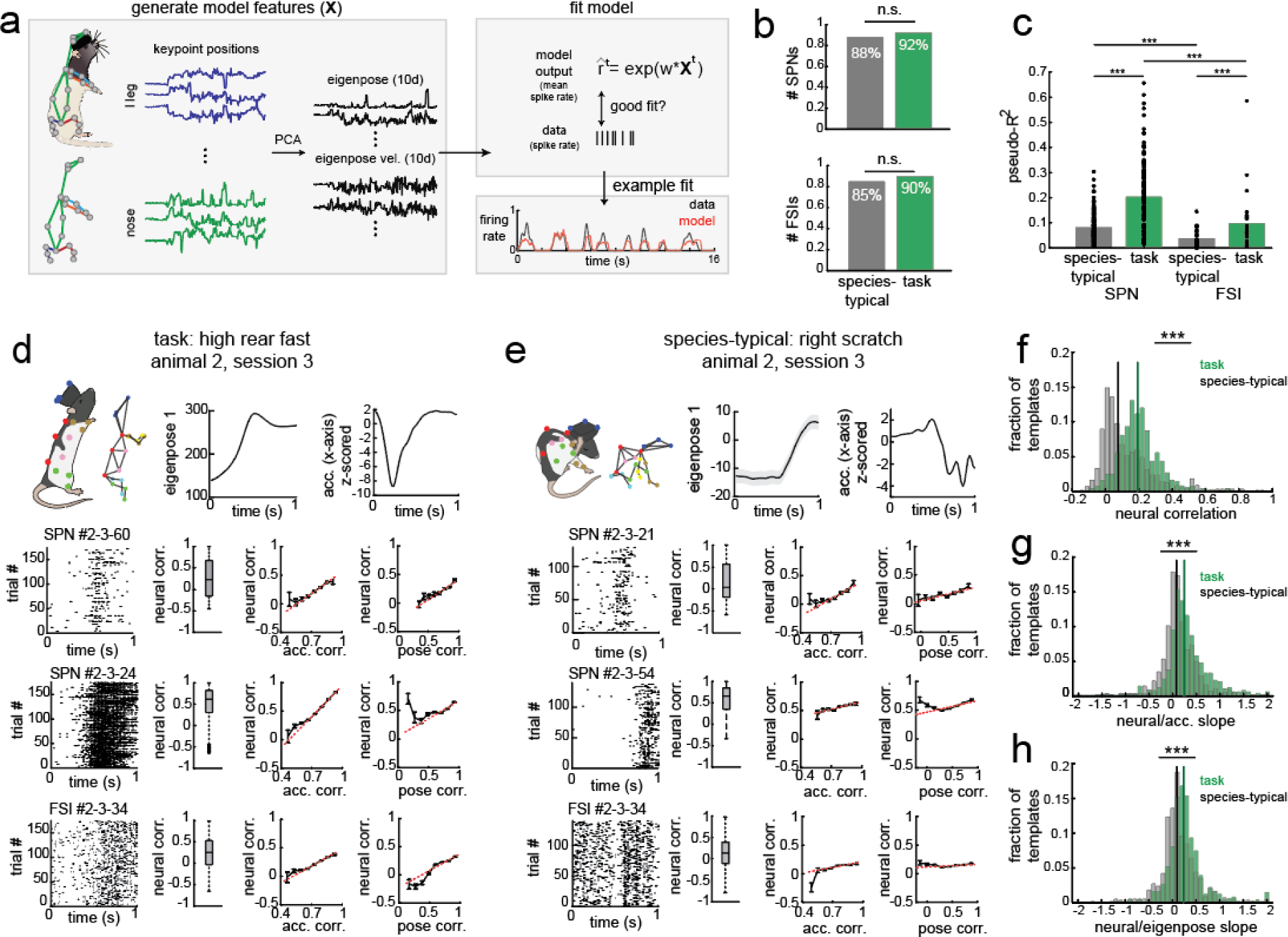
DLS neurons reflect movement kinematics across both behavioral domains but encode learned kinematics to a greater degree. **a.** Schematic of the GLM used to predict neural activity on held-out data. GLM features consisted of 10 eigenposes (marker data projected onto the top 10 PCs) and eigenpose velocity (the temporal difference of the eigenposes). Rat icons from scidraw.io, doi.org/10.5281/zenodo.3926125. **b.** The fraction of active SPNs (top) or FSIs (bottom) that encode kinematics, as quantified by the pseudo-R^2^ of each cell’s GLM, in each domain. There was no difference in the proportion of cells (SPN proportion test p = 0.21; FSI proportion test p = 0.57). **c.** The average pseudo-R^2^ for each cell on five-folds of held-out data for SPNs and FSIs with significantly high fits (pseudo-R^2^ > 0 across all five folds) during exploratory or task-related behavior. Task SPNs had a higher model fit than SPNs during exploratory behavior (ranksum p < 0.0001, task n = 127, species-typical n = 187), as did FSIs (ranksum p = 0.0087, task n = 26, species-typical n = 28). In both domains, FSIs had lower variance explained than the SPNs (species-typical: ranksum p < 0.0001, task: ranksum p < 0.0001). **d-e.** Top row: Schematic of the behavior (left), first dimension of eigenpose (middle) and the x-axis of the accelerometer (right) for matches corresponding to a task-specific template (d) or a species-typical template (e). Black line corresponds to mean signal across all template matches; gray corresponds to SEM. In some cases the gray line is hidden behind the black line. Bottom three rows: the spiking of example SPNs from during either template (left, cell numbers given in the title), a boxplot of the correlation in spiking across all trial pairs (middle-left), the relationship between correlation in the neural activity and correlation in the accelerometer signal (middle-right), and the relationship between the correlation in neural activity and the correlation in the eigenpose signal (far right). The red dashed line in the right two plots correspond to the best fit line across all data points. The error bars correspond to mean +/-SEM. **f.** The distribution of the average correlation in SPN activity across pairs of trials for task-specific templates (green) and species-typical templates (black). The average across-trial SPN correlation was lower for species-typical templates compared to task templates (task-specific template n = 543, species-typical template n = 481, ranksum p < 0.0001). **g.** The distribution of the slope between the average across-trial SPN activity correlation for each task-specific or species-typical template, where the across-trial SPN activity correlation is the average correlation in the firing rate across all possible pairs of trials, and the average across-trial accelerometer correlation, which is the mean correlation across accelerometer signals for all possible trial pairs. When averaging across neurons for each template, we observe that the slope is much higher for task-specific templates (ranksum p < 0.0001). **h.** The same as panel g, but for eigenpose signals instead of accelerometer signals. Again, we observe that the slope is much higher for task-specific templates (ranksum p < 0.0001).

Consistent with prior work^36^, we observed that the activity of nearly all active SPNs (127/138) and FSIs (26/29) were reliably modulated by the animal’s whole-body pose and movement during the timed lever-pressing task (Fig 5B). Interestingly, we observed that the fraction of active cells well-fit by the GLM during free exploration was similarly large (SPNs = 187/213, FSIs = 28/33). Indeed, there was no significant difference in the proportion of neurons encoding kinematics in the two domains (Fig 5B). Thus, if a DLS neuron is activated during a behavior, its spikes will correspond, at least coarsely, to the animal’s posture and movement, regardless of whether it is learned or species-typical.

However, even though just as many neurons encode kinematics, the kinematic code during species-typical behaviors could be more ‘muted’, i.e., reflect movements to a lesser degree, than during task-specific movements. To examine the strength of this code, we considered how well posture and movement could predict neural activity in the GLM for neurons that were sufficiently well-fit. We quantified model performance using the pseudo-R^2^ metric, where 0 corresponds to no extra variance explained by the model above assuming a constant average firing rate, and 1 is the maximum variance explained by a fully-saturated model^36^ (see Methods). While the model performed with similar predictivity as previously seen during the task^36^, with nearly 20% of spiking variance explained, its performance was much lower during species-typical behavior for both SPNs and FSIs (Fig 5C). This indicates that neural activity reflects movement to a greater degree during the task. However, as the distribution of behaviors differs between exploratory and task domains (Fig 4C), lower model performance during exploratory species-typical behavior could in fact stem from more variability in the input to the GLM.

To thus compare the kinematic code on more equal footing, we performed a second analysis for which variability across ‘trials’ could be more closely matched. Similar to our approach for analyzing the effects of lesions (Fig 3), we employed a template-based analysis to identify stereotyped repetitive movement patterns across behavioral domains and compared the associated neural activity. As in the previous analyses, we identified movement templates from the most frequently expressed finely-parsed behaviors (see Methods, n = 481 task templates, n = 543 species-typical templates). Templates were defined by the top 5 eigenposes, as well as from a 3-axis accelerometer that was affixed to the electrophysiology headstage, which further enhanced the identification of similar movements (see Methods for more detail).

Prior work has shown that most neurons active during the timed lever-pressing task exhibit precise firing patterns locked to kinematics (Fig S9, see also prior work^36^ for examples). Consistent with this, we observed that SPNs and, to some extent, FSIs, showed precise alignment to the task-specific templates (Fig 5D, Fig S10). We next examined whether this was also observed for templates identified during exploratory behavior. In line with our prior results, we found that both SPNs and FSIs exhibited behavior-locked spiking when aligned to matches of a species-typical template (Fig 5E, Fig S10). To probe whether there were quantitative differences in the precision or reliability with which spiking patterns reflected movements across the behavioral domains, we computed the mean correlation in the firing patterns across each pair of trials for each template (boxplots in Fig 5D-E). Consistent with neural activity reflecting task-relevant kinematics to a greater extent, the mean correlation of both SPNs and FSIs was substantially higher for the task-related templates (Fig 5F). Importantly, as species-typical template matches were also overall less stereotyped than for the task (Fig S10), this difference persisted even after accounting for any residual differences in trial-by-trial kinematic variability across domains (Fig S11A-H).

If DLS neurons indeed reflect the kinematics of task-relevant movement to a larger extent than for species-typical ones, as the above analysis suggests, then trial-to-trial variations in kinematics should, on average, also be more reflected in neural activity. To probe this, we computed the mean correlation across trial pairs for the kinematic features - for both the eigenpose and accelerometer - and examined the relationship between the variability in kinematics and variability in spiking (Fig 5D-E, right-most panels). This relationship can be quantified for each cell by computing the slope of the correlation in kinematic features versus the correlation in neural activity for all possible trial pairs. When comparing the average slope across all cells for each template in each domain, we observed overall steeper slopes, meaning stronger relationships, between movement kinematics and neural activity during the task for both SPNs (Fig 5G-H) and FSIs (Fig S11J-K). Again, this effect was preserved for SPNs when accounting for differences in the variability of behavior, although notably, this result did not hold for FSIs (Fig S11L-S).

### DLS codes for kinematics in behavior domain specific ways

So far, we have looked at statistics of firing and degree of kinematic tuning of DLS neurons and found significant, if moderate, differences across species-typical and task-related behavioral domains (Fig 4-5). However, our lesion results revealed a stark categorical difference in DLS’s contributions. Though it is plausible that downstream non-linearities could amplify the small differences in neural activity patterns to yield large functional differences, we considered an alternative, namely that the mapping between neural activity in DLS and kinematics could be very different across the domains^74^. Such a difference could explain the functional differences we see if one kinematic code (the one during skilled behavior) is far more effective in driving changes in downstream dynamics than the other^69^.

To address this possibility, we compared how DLS neurons represented movements across the behavioral domains, considering first the kinematic representations learned from the GLM in each behavioral domain (Fig 6A). By focusing on SPNs well-fit by models in both domains (n = 89 cells), we compared the model-predicted activity across various eigenpose values, as generated by parameter fits for the same cell and feature set. This revealed substantial differences in the kinematic encoding across species-typical and task-specific behaviors (Fig 6B), an indication that there may indeed be differences in the kinematic code across domains.

**Figure 6.**
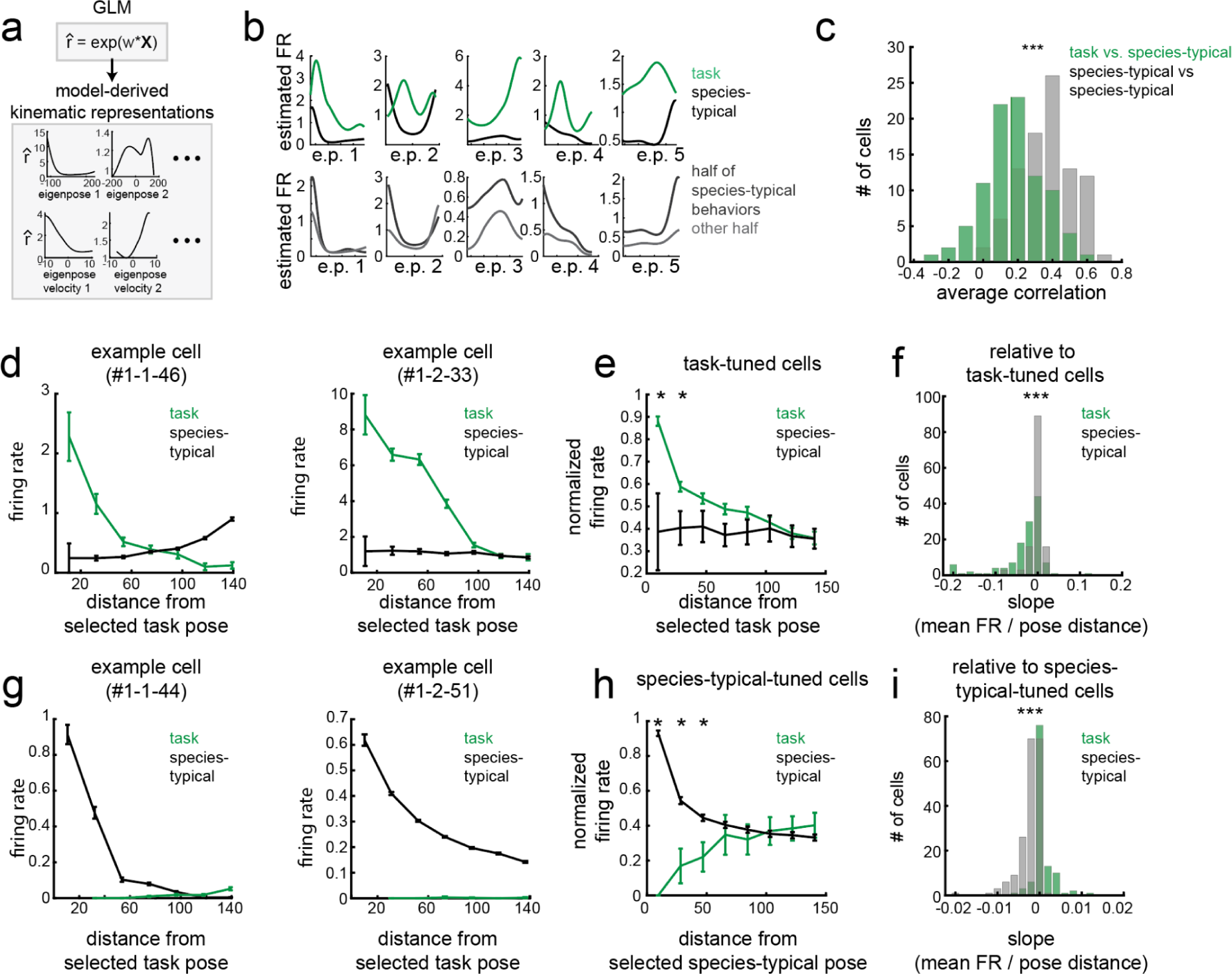
DLS neurons represent kinematics differently across behavioral domains. **A.** Schematic showing the model-derived kinematic representations (i.e. the estimated firing rates as a function of eigenpose and eigenpose velocity). These are computed using the learned parameters (*w*) for each cell’s GLM (see Methods for more detail). **B.** Model-derived kinematic representations for an example cell that encoded pose and pose velocity during both task and spontaneous behavior. Top row: model-derived kinematic representations computed from task or spontaneous behavioral domains. Bottom row: model-derived kinematic representations computed across two distinct subsets of species-typical behaviors. **C.** Histogram of the average correlation coefficient between model-derived kinematic representations in each behavioral domain (i.e. the correlation between data in the top row of B, in green), or across the distinct subsets of species-typical behaviors (i.e. the correlation between data in the bottom row of B, in black). The correlation was significantly lower across behavioral domains (n = 91, signed-rank test p < 0.0001). **D.** The average firing rate in each behavioral domain of two example cells as a function of the dissimilarity (quantified via Euclidean distance) from a pose selected from the task domain. The selected task pose is one that induces the highest activity for each cell. The mean activity of the same neuron is then analyzed for similar poses and distances during species-typical behavior. **E.** The average normalized firing rate (divided by the maximum value of the task-derived data, mean +/-SEM) as a function of the Euclidean distance from the selected pose. Only cells that were task-tuned, as determined through the GLMs, are plotted (n = 127). The stars above each bin correspond to significant differences (paired t-test p < 0.05/8, p-value corrected for multiple comparisons across bins; n data points for each bin for species-typical domain = 50, 93, 124, 127, 127, 127, 127, 127). **F.** The histogram of the slope values of the firing rate-by-distance curves that comprise the data in E. The slope of the cells during the task domain are comparatively lower than in the species-typical behavioral domain (signed rank test p < 0.0001, n = 127). The slope of the same cells when evaluated in the species-typical domain is not significantly different from 0 (signed rank test p = 0.77, n = 127). **G.** Similar to D, but the selected poses are computed from the species-typical behavioral domain. **H.** Similar to E, but for species-typical tuned cells, with the selected pose from the species-typical domain (as in G, n = 187). As in E, the stars above each bin correspond to significant differences (paired t-test p < 0.05/8, p-value corrected for multiple comparisons across bins; n for each bin: 4, 28, 53, 87, 113, 139, 160, 171). **I.** The slope for activity in the species-typical domain is comparatively lower than in the task domain (signed rank test p < 0.0001, n = 187). The slope of the same cells when evaluated in the task domain is greater than 0 (signed rank test p = 0.0002, n = 113).

To assess whether these differences were because the behaviors expressed during the domains were also different, we performed a control analysis, in which we split the set of species-typical behaviors in half and analyzed the parameters learned for GLMs fit separately to each half. One subset comprised behaviors similar to those expressed in the task, such as species-typical rearing that was similar to the rearing during lever-pressing, while the other subset included behaviors that were highly distinct from those in the task, such as grooming and scratching (see Methods). If differences in the kinematic representations across domains were driven by differences in the behavioral repertoire, the GLMs for the two subsets of species-typical behaviors should also be dissimilar. This, however, was not the case: the representations were remarkably consistent, suggesting a robust kinematic code for highly diverse behaviors (Fig 6C, Fig S12A). In comparison, there was a stark difference in the kinematic representation across behavioral domains, suggesting that the code is domain specific. The correlation coefficients between the kinematic representations across domains were much lower than between different subsets of species-typical behaviors. Similar results were also obtained when considering FSIs (Fig S12B-D).

We next verified this result using a model-free analysis, in which we examined the activity of the same neurons when the animal expressed similar behaviors across the two behavioral domains. Given the limited overlap in behaviors and that posture can explain more spiking variance than velocity (Fig S12E, also observed during the lever-pressing task in prior work^36^), we considered instantaneous pose as a proxy for kinematics. To compare pose-related activity across domains, we first selected the pose in a single domain that elicited the highest spiking activity for a kinematically-tuned cell (as detected by the model fit of its GLM). Note that we performed this analysis in a domain-specific way, i.e., a cell tuned to task-related behaviors had a preferred task-related pose. We then computed postural tuning curves relative to this ‘selected pose’ by computing the average firing rate for each cell as a function of the Euclidean distance between its selected pose and other poses observed in the same behavioral domain. Notably, within a behavioral domain, this curve has a negative slope, consistent with the kinematic code described thus far (Fig 6D-E, G-H).

If the kinematic code is the same across domains, a cell preferring a given pose in one domain should prefer the same pose in the other. However, we observed a striking departure from this: cells with a preferred pose in one domain would often display reduced activity for similar poses in the other domain (Fig 6D-I). We observed this both when taking the task domain, and task-tuned cells as a reference (Fig 6D-F), and when taking the species-typical domain, and species-typical-tuned cells as a reference (Fig 6G-I). In both cases, the magnitude of the slope between activity and postural distance was substantially lower and not significantly different from zero (Fig 6E,H). We observed similar trends in FSIs (Fig S12F-I), indicating the domain-specificity of the kinematic code is consistent across cell types.

Since this analysis focused solely on the instantaneous pose, we also considered sequences of similar poses. For example, it is possible that neurons exhibit different activity patterns when the same pose is expressed during different movements, such as a mid-rear pose expressed during a rear up or a rear down. We thus repeated the analysis using consecutive poses, with similar results (Fig S12J-M). Taken together, our results indicate that DLS represents the movement kinematics of species-typical and learned movements in strikingly different ways reflecting the profound difference in its functional contribution to movement kinematics in the two domains.

## Discussion

Our study considers a fundamental distinction in how the BG contribute to movement, specifically asking whether this region is essential for general control or specialized for acquiring and controlling learned motor skills (Fig 1). Using high-resolution 3D kinematic tracking in freely behaving rats, we showed that lesions of the sensorimotor striatum did not affect the expression (Fig 2), kinematics (Fig 3) or sequencing (Fig S6) of naturally expressed species-typical behaviors. This was in stark contrast to learned task-specific movements, which were profoundly affected by the same lesions (Fig 3, see also prior work^36,41^). To probe the neurophysiological correlates of this difference, we recorded from the same cells in sensorimotor striatum during both learned and species-typical behaviors. We found a small but significant difference in the firing statistics across the domains (Fig 4) and a stronger kinematic encoding of learned behaviors (Fig 5). Importantly, while striatum reflected ongoing movements at all times, the mapping between neural activity and motor output was dramatically different across behavioral domains (Fig 6). Taken together, these results suggest that the basal ganglia are specialized to control learned movement kinematics, and may do so by learning task-specific kinematic codes that effectively and adaptively influence dynamics in downstream control circuits.

### BG’s role in basic motor control

Our lesion results, which suggest that the BG are not essential for basic movement control, are at odds with established models informed by studies of BG disorders^78^, many of which result in severe control deficits^25^. As unnatural alterations of BG dynamics in both humans and animal models can affect basic control, it has been argued that these disruptive effects reflect ‘gain-of-abberant-function’ rather than an obligatory role for the BG in the affected control functions ^4,31^. In Parkinson’s disease, for example, imbalances in direct and indirect pathway activation can lead to a pathological rhythmicity in BG’s output^79–81^, the behavioral effects of which can be ameliorated by suppressing or regularizing the activity through lesions or stimulation^35,82,83^. Consistent with this, mimicking the pathophysiology of Parkinson’s disease in rodents by inducing degeneration in the nigrostriatal pathway results in far more severe control deficits than what is seen after DLS lesions^78,84^.

Yet prior studies in rodents have reported deficits in species-typical behaviors also after DLS lesions. These include the premature termination of grooming sequences^61,62^ and changes in the expression and sequencing of other types of naturalistic behaviors^28^. Despite inducing large bilateral DLS lesions and recording naturalistic behaviors at unprecedented resolution, we could not replicate the high-level sequencing effects (Fig S6), nor did we see any changes to the fine-grained kinematics of species-typical behaviors (Fig 2-3).

A methodological difference between ours and previous studies could potentially account for these discrepancies. In prior studies showing lesion-induced effects on species-typical behaviors, DLS in both hemispheres was lesioned at the same time by overstimulating its neurons with excitotoxic agents. Such overstimulation can induce lasting effects on downstream brain circuits^85–87^, thus potentially altering their function in both the short and long-term. The severity of these off-target effects is a function of lesion size, and hence can be ameliorated by performing them serially^56,88^, which is what we did in this study by lesioning one hemisphere at a time (see Methods). That we do not see any effects suggests that the BG are not part of the essential minimal circuitry required to express species-typical behavior in an exploratory context.

### Differential effects of striatal dynamics on behavior

The lack of behavioral effects of DLS lesions on species-typical contrasted with dramatic effects on learned movement patterns. This suggests that the BG is differentially engaged across different types of behaviors and can ‘switch’ between shaping the detailed motor output for learned movements while taking a back seat for behaviors adequately generated by downstream control circuits, i.e. species-typical behaviors. How a circuit, like the BG, switches between effective and ineffective ‘modes’ is an intriguing question that pertains to most flexible and modular systems^69,89^.

For example, while both motor cortex and basal ganglia are integral parts of the distributed mammalian control system and *can* drive motor output, it does not follow that they always should or, indeed, do. Consider normal locomotion, a repetitive behavior implemented largely by specialized circuits in the brainstem and spinal cord^90^. While both areas are active during locomotion^28,69,91^, neither is required for generating the repetitive movement patterns^39,69^. This has prompted the idea that neural activity are, in such circumstances, in the ‘null-space’ for brainstem- and spinal cord-generated behaviors, i.e., aligned with dynamics in the downstream control circuits in such a way that makes the activity ineffective in driving changes^69,92–94^. Since the ‘null-space’ for a circuit likely depends on the state of the downstream control regions, the activity of its neurons should reflect ongoing behavior (i.e. state changes) whether it contributes to control or not^69,95^. Our results are consistent with such a picture.

While we find that DLS encodes movement at all times (Fig 4-5), the mapping between neural activity and kinematics (Fig 6) depends critically on whether the behavior is learned or species-typical. Thus, a possible mechanistic explanation for why the BG take a back seat during species-typical behaviors is that their activity in this domain is in the motor ‘null-space’. The process of learning could induce changes to BG dynamics, shifting their output into a motor-potent sub-space, thus making the BG an essential part of the control circuit by driving brainstem controllers. Acute manipulations of BG activity and imbalances in network dynamics brought on by disease (e.g., Parkinsons) could similarly shift BG’s output into motor-potent subspaces, including during exploratory movements. In these cases, however, the abnormal activity patterns would compromise, not aid, behavior.

Firing statistics, specifically the degree of burstiness^96,97^, is another aspect that could influence the degree to which BG affects movement. Notably, basal ganglia pathologies that affect normal movement control are often associated with increases in burstiness and rhythmicity^98,99^. That we see DLS activity trending in this direction also during learned behaviors (Fig 4) could help explain the functional differences across behavioral domains, especially under the assumption that these differences are then further amplified through nonlinearities in the BG circuit^68^.

### The role of BG in learning

While we did not find evidence for the DLS controlling species-typical behaviors, our analysis of motor skills suggested, in concert with earlier studies^36,38,39,41^, that it is essential for shaping learned movement kinematics. Densely innervated by dopaminergic neurons signaling reward prediction error^100^, sensorimotor striatum could learn to map state input from cortex and thalamus to activity patterns that help shape future actions in adaptive and task-specific ways^2,101^. The kinematic representation in DLS during species-typical behaviors, albeit ineffective for influencing movements, could thus serve as a scaffold for learning.

While BG’s role in motor learning is well established, their role in controlling learned movement patterns does not automatically follow. For example, in songbirds, the song-specialized BG (Area X) ‘tutors’ cortical circuits during learning^4,102^, ‘handing off’ what it learns to nucleus robust archopallium (RA), a motor cortex analog brain area dedicated to controlling the courtship song. However, many learned behaviors, including the movement patterns we analyzed in this paper, are generated by brainstem controllers. Modifying these circuits for a given task would have the effect of altering and interfering with the many behaviors and control functions they subserve.

Instead, our findings suggest that the BG may be part of a circuit, that likely also includes thalamus and brainstem^11,103^, that stores learned policies and helps control skilled movements by interacting with more hardwired brainstem circuits in task-specific ways. This, of course, does not preclude BG also ‘tutoring’ cortex, transferring learned aspects of behavioral control to cortical circuits, which - unlike brainstem circuits - have the redundancy and capacity to adapt in task-specific ways without unduly interfering with other behaviors^104^. How the learning process by which the BG assumes control over motor output plays out, and indeed, how activity patterns and coding schemes in DLS change with learning from passively reflecting motor output to activity driving remains to be understood.

## Methods

### Ethics Declaration

The care and experimental manipulation of all animals was reviewed and approved by the Institutional Animal Care and Use Committee at Harvard University Faculty of Arts and Sciences.

### Animals

Subjects for all experiments were Long-Evans rats (Charles-Rivers Laboratories). For the lesion experiments, we used female rats aged 3-5 months at the start of the experiments (n = 4 bilateral DLS lesions, n = 4 sham lesions). For the electrophysiology recordings, we used bothmale or female rats, aged 2-6 months at the start of the experiments (n= x male n = x female). No statistical methods were used to predetermine the number of subjects in our study.

### Surgical Procedures

All surgical procedures were designed to limit pain and discomfort and were performed under 1-2% isoflurane anesthesia. Prior to surgery, all tools were sterilized using a hot bead sterilizer and rinsed in 70% ethanol. Animals received painkillers (buprenorphine or carprofen) after surgery and were allowed adequate recovery time before proceeding with any behavioral experiments.

### Lesion surgeries

Bilateral lesions of the DLS were performed in a two-stage procedure following previously established methods^36,57^. Animals were anesthetized with 2% isoflurane in carbogen and securely positioned in a stereotactic frame. A midline incision was made on the scalp to expose the skull, and Bregma was identified as the reference point. Four small craniotomies were created above DLS for injections of quinolinic acid (0.09 M in PBS (pH 7.3)) or sterile saline. The following coordinates were used for each injection (relatove to Bregma and midline): (AP +0.7, ML +3.6, DV +4.8), (AP +0.7, ML +3.6, DV +4.1), (AP −0.3, ML +4.0, DV +5.1), and (AP −0.3, ML +4.0, DV +4.1). Quionolinic acid, an excitotoxin, was administered using a fine glass pipette, attached to a microinjector, which was lowered to the desired injection site. Each injection consisted of 35 pulses of 4.5 nL each. To facilitate diffusion and prevent backflow of the drug, the glass pipette was slowly retracted after each injection. Once all injections were completed, the incision was sutured, and the animals received painkillers to minimize discomfort during recovery. After the first stage of lesioning, animals were allowed to recover for 10-14 days before undergoing a similar lesion on the contralateral hemisphere. Following the second lesion, animals were given another 14-day recovery period before resuming behavioral recording.

### Tetrode implant surgeries

Prior to surgery, we constructed microdrives containing individual tetrodes as previously described ^46^. In brief, we twisted strands of 12.5 micron nichrome wire and threaded them through an array of seven 34-gauge polyimide tubes mounted on a custom microdrive. One end of the tetrodes were secured to a custom interface board, digitally amplified at 16-bit resolution using a 64-channel headstage (Intan Technologies). We electroplated the tetrodes in a gold and carbon nanotube solution to achieve an average electrode impedance of 100-150 kOhm.

We implanted microdrives into DLS following established protocols ^36^. We removed hair on the scalp, affixed the animal to a stereotaxic apparatus, and sterilized the scalp with betadine and 70% ethanol. A longitudinal incision was made and skin was carefully removed to expose the skull. Five skull screws were inserted at midline, over the cerebellum, visual cortices, and frontal cortices. A 200 um silver ground wire was attached to the cerebellar skull screw to serve as a grounding reference, while a 50 um steel reference wire was inserted as a reference electrode. A 4 mm diameter craniotomy, centered over the DLS at coordinates (+4.25 mm ML, +0 mm AP), was performed. After removing the dura, the tetrode bundle electrodes were slowly inserted to a depth of −4 DV mm. The tetrode drive, along with a protective headcap, was securely attached to the skull using Metabond and dental cement. To optimize recording quality, adjustments to the tetrode depth were made at one-week and two-week intervals post-surgery.

#### Motion capture marker attachment surgeries

Marker attachment surgeries were performed following the tetrode implant and followed previously described protocols ^44^. Markers were fabricated using 5 mm diameter high-index of refraction ball lenses ^44^ (N=2.0, H-ZLAF, Worldhawk Optoelectronics). We used high-strength epoxy to affix retroreflectors to steel earrings that we attached to the limbs or to a monel clamp attached to the trunk. Three retroreflectors were attached to a 3D printed acrylic headcap painted using flat black paint to track the head.

To prepare for the marker attachment, body piercings were sterilized using 70% ethanol, and the animal’s head, trunk, and limbs were shaved. Marked sites for the body piercings were designated using a skin pen, and the marked areas were sterilized with alternating washes of betadine and 70% ethanol. For markers on the spine, trunk, and hips, two small incisions spaced 1 cm apart were made, and the body piercings were inserted through the incisions and fastened using pliers. For markers on the shoulders, forelimbs, and hindlimbs, similar procedures were followed, but with incisions spaced 10 mm apart. To secure the limb piercings, earnuts were attached to the back of the piercings. Damped wooden barriers were used to create space between the earnuts and the skin, and a soldering iron with flux was used to solder the earnuts to the piercings. We applied antibiotic cream to the marker sites and administered subcutaneous injections of buprenorphine (0.05 mg/kg) and carprofen (5 mg/kg) following the surgery.

### Data collection

#### Video recordings of DLS and sham-lesioned animals for pose tracking

Video recordings of both DLS and sham-lesioned animals were obtained to track their natural behavior in a 2-foot diameter plexiglass cylinder. We recorded each animal for 1.5 hours each day for 4 days before the first surgery and again for four days two weeks after the second surgery. Animals were familiarized with the arena prior to the first recording. Recordings were performed at similar times (+/-1 hour) each day. Video frames (1,320 x 1,048 pixels) were sampled at 50 Hz from 6 synchronized cameras (Flea3 FL3-U3-13S2C). One camera, which was miscalibrated, was excluded from the analysis.

Before the initial recording, the cameras were calibrated using custom MATLAB scripts that utilized the Camera Calibration functions from the Computer Vision toolbox. Intrinsic camera properties were determined using a black and white checkerboard pattern, and extrinsic camera parameters were computed by manually marking four points on an L-shaped frame. Similar calibration procedures were conducted after the final recording session. Animals were habituated to the recording environment for several days before the first session, and they were kept on a normal 12-hour light/dark cycle with access to ad libitum water and food in between behavioral recording sessions.

#### Electrophysiology and CAPTURE recordings

Motion capture recordings were conducted using a 12-camera motion capture system (Kestrel, Motion Analysis). The cameras were positioned 5-10 feet from the center of the 2-foot diameter plexiglass cylinder, which was the same environment used for recording the lesioned animals. The cameras were elevated to two different heights and angled at 15° and 35° to the horizontal. A custom lever and water spout were installed in the arena for animal training, and the arena was kept on a 12-hour light cycle.

Prior to all recordings, rats were previously trained on the timed lever-pressing task following established training paradigms ^36,57,105^ in automated behavioral training boxes. Animals were water restricted and given three one-hour training sessions per day. During these sessions, the rats were required to press a lever twice within a 700 ms interval in order to receive a water reward. Failed trials were penalized with a timeout of 1.2 seconds. The rats were pre-trained until they achieved a mean inter-press interval within 10% of 700 ms and a coefficient of variation of less than 0.25. Following pre-training, these rats were habituated to the motion capture arena until consistent and reliable task performance was observed. Recording sessions in which the number of lever taps fell below the threshold of <150 were excluded from analysis. The excluded sessions mainly consisted of early exposures to the CAPTURE environment following initial training.

Electrical activity of cells was recorded at a sampling rate of 30 kHz and digitized using a custom-built cable connected to an FPGA (XEM6010, Opal Kelly), which in turn connected to a Windows machine running custom electrophysiology acquisition software ^46^. The data was acquired continuously and subsequently divided into distinct ‘sessions’ based on the day of recording.

### Histology

After completion of behavioral and electrophysiological recordings, animals were euthanized with a mixture of ketamine (100 mg/kg) and xylazine (10 mg/kg). To verify tetrode placement, an anodal current was passed through select electrodes to create microlesions at the tip. Subsequently, animals were transcardially perfused with 4% paraformaldehyde. Their brains were harvested for histological examination to confirm either lesion size and location or the targeting accuracy of the implanted tetrodes. In both cases, the brains were sectioned into 80µm slices using a vibratome, mounted onto glass slides, and stained with cresyl violet. The stained slices were imaged using a slide scanner.

#### Quantification of lesion size

For accurate estimation of the total lesion size, we analyzed seven sections spanning the anterior-posterior extent of the striatum (Bregma +1.70, +1.20, +0.70, +0.30, −0.80, −1.30 mm). Lesion boundaries were manually marked using Adobe Illustrator, based on differences in cell morphology and density, such as the loss of larger neuronal somata and accumulation of smaller glial cells. The striatal boundaries for DLS and DMS were determined based on prior work ^36^. Neighboring brain regions were identified using the Paxinos Rat Brain Atlas (4th edition) ^106^ using anatomical landmarks (external capsule, ventricle) and cell morphology and density. The fraction of each lesioned area was quantified from images in Fig S2A. In all animals, at least 50% of DLS was lesioned in both hemispheres (Fig S2B).

#### Verification of accurate tetrode placement

For confirming precise tetrode placement, we analyzed sections spanning the anterior-posterior extent of the striatum (Bregma +1.70 to −1.30 mm). Striatal boundaries for DLS were determined based on prior work ^36^ and mapped onto sections with clear microlesions using Paxinos Rat Brain Atlas (4^th^ edition) ^106^. Tetrode tip location was then marked by a black arrow (Fig S6). The tetrode tip locations were then marked with a black arrow (Fig S6). All recordings successfully targeted DLS, although the final recording location varied along the dorsal-ventral axis.

### Data processing

#### Markerless pose tracking using DANNCE

Pose estimation with DANNCE involved two main steps: center of mass (CoM) detection and DANNCE keypoint estimation. Both steps were implemented in Python 3.7.9, utilizing standard packages for scientific computing and deep learning (following ^43^). The code and initial weights for pre-trained networks were obtained from the Github repository (http://github.com/sponsso/dannce/).

To train the 3-D U-Net for detecting the center of mass of the animal in the frame (the ‘CoM network’), we used 1000 hand-labeled frames from all animals before and after surgery. The hand-labels were generated using Label3D, a Matlab-based graphical user interface specifically designed for generating 3D hand-labeled poses. The CoM networks were trained as previously described in ^43^.

Once the CoM network reliably detected the center of the animal in novel frames, we labeled a 20-keypoint skeleton. We manually labeled the 3D positions of approximately 1600 frames across sessions from all animals involved in the lesion experiments before and after surgery (n = 8 animals) using Label3D. For fine-tuning this model, we followed the approach of prior work, finetuning a model previously trained to track keypoints in the Rat7M dataset on our training set. Computational resources in the Cannon High-Performance Cluster operated by Harvard Research Computing were utilized for this purpose. To assess the labeling accuracy, we compared DANNCE labels with the output of two expert human labelers, evaluating all 20 keypoints over 80 frames collected across all 8 animals.

#### Motion Capture Processing

Preprocessing of motion capture data was conducted on a custom analysis workstation (128 GB RAM, 3.6 GHz Intel i7) using Matlab (Mathworks, Natick MA). We applied a 3-frame median filter to smooth marker data and placed all markers in a reference frame centered on the middle of the animal’s spine, with the front of the spine oriented on the y-axis. Markers were designated as missing if they were not detected, fit by the motion capture body model, or exceeded a velocity threshold of 25 mm/frame. In addition, markers on the forelimbs and hindlimbs showing non-physical configuration relative to markers on the olecranon or patella were also considered missing.

#### Spike sorting and unit identification

Recordings were analyzed using the Fast Automated Spike Sorter (FAST) as previously described^46^. FAST automatically generated a candidate set of units, which were manually inspected and categorized as accepted, merged, rejected, or split using a custom MATLAB graphical user interface. Units were rejected if they had an amplitude <75 μV, were identified for less than 5 hours, displayed a high percentage of inter-spike intervals <1 ms, or exhibited waveforms consistent with motion or electrical artifacts. Units identified across day-long recording sessions were merged based on the similarity of their waveforms. In some cases, automatically identified units were split using mClust4.2 based on waveform amplitude, shape, and energy when they consisted of two units with distinct waveforms.

Isolated units were classified as putative SPN (striatal projection neuron) or FSI (fast-spiking interneuron) types based on spike-waveform features, including peak width (full width at half maximum) and time interval between spike peak and valley, as well as their mean firing rates averaged over the recording duration of each unit. Units meeting the criteria of peak width >150 µs, peak-valley interval >500 µs, and mean firing rate ≤10 Hz were classified as SPNs, while units with peak width ≤150 µs, peak-valley interval ≤500 µs, and mean firing rate ≥0.1 Hz were classified as FSIs. Units not meeting any of these criteria were excluded from all analyses. If a unit didn’t exhibit spikes for a continuous 10-minute period, we omitted this data from analysis due to the possibility of spontaneous drop-out.

### Data analysis

*Applying ‘MotionMapper’ pipeline to DANNCE-generated 3D keypoint data: Lesion dataset* In order to identify and cluster distinct behaviors using the keypoint data, we first reduced the high-dimensional keypoint data for all sessions to a 2-dimensional space, smoothed it, and then applied watershed clustering. This process follows variants of procedures described in prior work^44,48,107^.

To embed behavioral data into a low-dimensional space in a way that promotes the clustering of distinct behaviors, we first computed a ‘feature vector’ for each timepoint of each behavioral session. Each feature vector was composed of three feature types: a wavelet decomposition of the egocentrically-aligned keypoint data projected onto the top 10 principal components, the velocity of each keypoint, and the z-value (height) of each keypoint. Egocentrically-aligned data was based on two keypoints on the animal’s back, where one defined the origin and the other indicated the direction. The wavelet decomposition considered 25 frequencies, ranging from 0.25 to 20 Hz, with power among these frequencies indicating the rate of change in the animal’s posture over time. To ensure equal weighting of signal types, we applied logarithmic transformations to the watershed signal and considered values below a threshold (−8). Additionally, we multiplied the z-value by 0.1.

Given the unbalanced nature of the data, with certain behaviors occurring more frequently than others, we employed importance sampling. This method allowed us to create a representative behavioral sample set that appropriately accounted for rare behaviors. The process involved using t-distributed stochastic neighbor embedding (t-SNE) to reduce the set of feature vectors for each session into a 2-dimensional space. We then selected a set of 1000 distinct points (feature vectors) from this reduced plot for further analysis. After applying this process across all sessions, we combined the sub-selected feature vectors, totaling 64,000 vectors, and reduced them into a 2-dimensional space using t-SNE. This space served as the behavioral map into which all feature vectors from all sessions were embedded, such that feature vectors with similar values were placed in proximity on the map.

To identify distinct behaviors from this map, we next performed watershed clustering. We binned the map into 500 x 500 bins, computed the number of points in each bin, smoothed it using a Gaussian filter (sigma = 1.5 bins), and then performed watershed clustering to achieve 50-80 distinct behaviors. Each behavior cluster was visually verified by inspecting sample video snippets, after which they were manually labeled. These labels facilitated the aggregation of behaviors into 16 broader categories.

Lastly, to improve the assignment of each time point to a fine or coarse cluster, we identified fast and slow movements across the behavioral space using a Gaussian mixture model. As fast movements across behavioral space likely correspond to transitions between behaviors^48^, we imputed these values according to the starting and ending behavior. We considered only behaviors with a dwell time in a finely-parsed cluster of 10 time bins (200ms). Time points that did not satisfy this criteria were filled in with the nearest (temporally) cluster.

To enhance the precision of assigning time points to clusters, we first filtered the sequence of behavioral clusters over time. We retained only those periods during which the animal remained within a single finely-parsed cluster for 10 consecutive time bins. If any time points did not meet this condition, we substituted them with data from the nearest temporally-adjacent cluster

#### Jensen-Shannon divergence

To compare the distribution of finely-parsed behaviors, coarsely-parsed behaviors, and the distribution of points within each finely-parsed behavior, we employed the JS-divergence^108^. This is a symmetric quantity that quantifies the difference in two probability distributions P and Q, and is defined as: 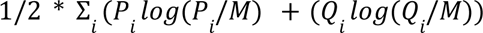, where *M* = 1/2 * (*P* + *Q*).

#### Comparing the distribution of feature vectors pre- and post-lesion

For each finely-parsed cluster, we first identified all instances when the animal occupied that particular cluster, then matched these instances with their corresponding pre-lesion and post-lesion feature vectors. In cases where consecutive instances occurred for a maximum of 10 time points, we calculated the average of the feature vectors. We then computed the average distance between all the averaged pre-lesion feature vectors (using the pdist2 function in MATLAB), the average distance between all pre-lesion and post-lesion feature vectors, and normalized the pre-post distance comparison by the pre-only comparison.

#### Comparing vigor of grooming and running speed

To compare grooming bouts before and after lesion, we considered each finely-parsed grooming cluster separately. We identified grooming bouts as instances where at least 30 consecutive seconds were spent in a grooming cluster. Given that some bouts included body grooming, we filtered bouts by examining arm movements. To do this, we calculated the grooming power, which is the power in the 2-6 Hz frequency band divided by the power in the 0-14 Hz band, for the 3D egocentric wrist positions within each bout (Fig S4). For each bout with a power above 0.3, we then also computed the number of grooming cycles by computing the number of peaks, as well as the amplitude (mean distance between each peak and trough) of each grooming cycle.

To assess whether these grooming metrics (grooming power, peak count, and amplitude) changed after surgery for each DLS-lesioned animal and grooming bout, we employed mixed-effects models. These models accommodate both random and fixed effects, which is crucial when dealing with hierarchical data structures (e.g., when multiple data points come from each animal). In our analysis, we treated each grooming metric as an observation (the dependent variable to be predicted for each model), the pre-or post-DLS lesion status as the fixed effect, and the identity of the animal and grooming cluster as random effects (returning 16 distinct data types, for four animals and four grooming types). We implemented these mixed-effects models using MATLAB’s ‘fitlme’ function, allowing us to assess whether the fixed effect had significant predictive power on the grooming metric.

To compare the mean speed, we computed the animal’s running speed at each time point using a single marker on the trunk. We then identified times of movement by applying a speed threshold of 5 cm/s, and computed the average running speed for each session.

#### Template-based analysis for the timed-lever pressing task

We re-analyzed data used in a previous publication^36^. In this study, the positions of both hands in two-dimensional space were tracked using DeeperCut as animals learned to perform the timed lever-pressing task. After task performance plateaued, animals received either a DMS (n = 5) or DLS (n = 6) bilateral lesion. To quantify a 1-second template in the task, the movement data from 0.15 seconds before the first tap to 0.85 seconds after the first tap were analyzed. The most common movement pattern expressed in the 500 trials with an inter-press interval (IPI) ranging from 650 to 750 ms was identified using hierarchical clustering. Clusters with a minimum correlation of 0.95 across the four signals (x and y coordinates of the left and right hand) were considered. The average of the largest cluster was taken as the template.

To compare expression of this template after lesion, the last 5000 trials (maximum) with an IPI between 650 and 750 ms were examined. The average correlation of each trial to the template was calculated, and the number of trials with an average correlation greater than 0.75 was determined. This was compared to the first 5000 trials after surgery with the same IPI range.

#### Template-based analysis for species-typical behavior

Grooming: For each grooming bout longer than 1 second prior to surgery, we identified 1-second bouts with at least two identifiable cycles and a grooming power (power in 2-6 Hz / power in 0-14 Hz) above 0.3 (following procedures described in “*Comparing vigor of grooming and running speed”)*. Bouts that were longer than 1 second were divided into 1-second snippets that were then separately analyzed. One second bouts were chosen, as longer snippets made it difficult to identify precise replicates, while shorter snippets would encapsulate only a cycle or two and provide an insufficient basis for comparison. From each selected grooming snippet we cross-correlated the 3D egocentric wrist position with other sessions from the same animal, both before and after the surgery, to search for matches with a correlation greater than 0.75^63^. Matches for a single template were greedily chosen to avoid overlap between matches. Snippets were recorded as templates if they had more than 5 matches in the pre-surgery dataset. To avoid redundancy, templates and their matches were removed if they overlapped with another template and its matches by more than 10%. In such cases, the template with the most matches was retained.

Whole body movements: For each animal, we initially identified their unique ‘eigenposes’ by projecting the flattened 3D egocentric keypoint data onto the top 5 principal components. We then detected instances when the animal engaged in ‘active’ movements, such as rearing, walking, or exploring (Fig 3). We then identified snippets as 1-second bouts within each active movement. Snippets were filtered based on their variance (such that flat snippets were ignored). For each snippet, we searched for candidate matches in the data both pre- and post-surgery by cross-correlating it with the eigenpose data from all sessions of the anima, recording all matches with a mean correlation coefficient > 0.75. In addition, we computed the average Euclidean distance between each template and candidate match, rejecting matches for which the distance exceeded at threshold (70). Similar to the grooming template identification process, a snippet was classified as a template if it had more than 5 matches pre-surgery. Overlapping templates and their matches were discarded.

Comparison of template matches using mixed-effects models: To compare the number of matches before and after surgery for each template in each animal, we employed mixed-effects models. This modeling approach naturally accommodates the hierarchical structure of our data, where each animal possesses a unique set of templates. In our analysis, the number of matches post-surgery divided by the number of matches pre-surgery for each template served as the observation (the dependent variable) in each model. We considered the type of lesion (sham or DLS lesioned) to be the fixed effect, while the identity of the animal was treated as a random effect. We implemented these mixed-effects models using MATLAB’s ‘fitlme’ function, enabling us to assess whether the lesion type had a significant predictive impact on the number of template matches observed after surgery.

#### Examining behavioral transitions

We computed transitions between finely-parsed or coarsely-parsed behaviors by computing the fraction of time behavior B followed behavior A, divided by the amount of time spent performing behavior A such that the rows of the transition matrix sum to 1. We ignored self-transitions (i.e., a behavioral string of 1, 1, 1, 2, 2, 3, became 1, 2, 3).

To determine whether the transition matrix changed more after DLS lesions compared to sham lesions, we computed the modulation in transition matrix values, defined as *P_modulation_*= (*P_pre_*− *P_post_*)/(*P_pre_* + *P_post_*), and compared the average absolute value of P*_modulation_* across cohorts.

To compute the entropy (*H*) of the transition matrix (*P*) before and after surgery, we following previously described procedures^28,109^: *H*(*P*) = Σ*_ij_* Π*_i_ P_ij_ log*(*P_ij_*), where Π*_i_* is the probability distribution for behavior *i*. We computed the entropy of behavioral transitions for finely-parsed behaviors, coarsely-parsed behaviors. To compute the entropy of the grooming transitions, we considered the four finely-parsed grooming behaviors, and grouped all other behaviors into a single ‘other behavior’ category.

#### Applying keypoint MoSeq to the behavioral data

To analyze whether changes in entropy could be determined through Keypoint MoSeq^67^, we downloaded the code from the github repository: https://github.com/dattalab/keypoint-moseq (downloaded June 2023). We rotationally aligned the data in a similar manner to our egocentrically-aligned data, using two keypoints on the trunk to determine the origin and direction. We used the default value as given in the code for the syllable duration parameter κ (κ = 90000). Under this model, we identified 17-33 frequently-used (as indicated by the code) syllables per animal (∼90 syllables total per animal).

#### Applying ‘MotionMapper’ pipeline to CAPTURE-generated 3D keypoint data: Electrophysiology dataset

Behaviors were clustered for the CAPTURE-generated dataset using a similar analysis pipeline as the DANNCE-generated dataset. First, the marker data was de-noised. Statistically aberrant marker positions were imputed. Aberrant marker positions were detected using the average distance between egocentrically-aligned markers (e.g. the distance from the snout to the ear should always be similar). Imputed marker positions were then smoothed with a median filter and downsampled from 300 Hz to 50 Hz. We then created a set of feature vectors for each time point using the same three feature types (wavelets of the egocentrically-aligned data, the marker velocities, and the height of each marker). We projected the set of feature vectors into a 2D space using t-SNE, created a training set to embed the data by subsampling a range of 6500 behaviors in 15 sessions. We then combined the subsampled feature vectors, projected those into a 2D space using t-SNE, and embedded all remaining feature vectors into this space. We then computed a smoothed density of points, performed watershed clustering, visually inspected each cluster, assigned each cluster behavioral label, and then groups labels to define coarsely-defined clusters. For most analyses involving the electrophysiology data, we excluded the ‘idle’ clusters.

For the majority of analyses, ‘task’ times were determined to be the time within 5 seconds of any lever press. The vast majority of lever presses occurred during defined sessions times. Times of exploration were times when the animal was actively behaving (i.e., not idle) and had not pressed a lever for at least 10 seconds.

#### Quantification of average firing rate

For each cell, we computed the average firing rate (# of spikes / # of time points) in each behavioral domain across coarsely-defined behaviors with at least 30 seconds of data.

#### Quantification of bursting

We computed the interspike interval (ISI) for each cell by taking the temporal difference between neighboring spikes (sampled at 300 Hz). Bursting was quantified by the fraction of ISIs less than 10 ms (corresponding to bursting > 100 Hz). Only cells that were ‘active’ (avg firing rate > 0.5) in either domain were considered in this comparison.

#### Quantification of correlation in firing rates between cell pairs

To correlate the firing rate of spike trains, we binned spike trains into 12 Hz (∼80 ms) bins, smoothed the firing rate with a Gaussian filter with width = 10 bins, and then correlated the smoothed firing rate for each pair of cells. Only cells that were ‘active’ (avg firing rate > 0.5) in either domain were considered in this comparison.

#### Generalized linear model (GLM)

GLMs, also referred to as a linear-non-linear Poisson (LN) model, were fit to each neuron to determine if spiking could be predicted as a function of behavioral features. Specifically, these models quantify the dependence of spiking on a set of behavioral features by estimating the spike rate 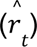 during time bin *t* as an exponential function of a sum of inputs, which here include a baseline firing rate (*b_o_*) and a time-varying input derived from the learned contribution of each behavioral feature at time *t*. For each behavioral feature, this input is computed as the dot product of the feature vector, which denotes the specific values of the behavioral feature at time *t*, with a corresponding set of parameters learned by the model: 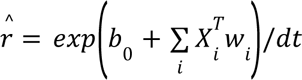. Here *i* indexes the variable, *X_i_* is a *W_i_ x T* matrix where each column is a behavioral feature vector (*x*_*i*_ length *W_i_*), *W_i_* is a *W_i_ x* 1 column vector of learned parameters that converts behavioral feature vectors into a firing rate contribution, and *dt* is the time bin length (83 ms for 12 Hz sampling).

For the behavioral features, we used 20 spline functions^110,111^ relating to the eigenpose and eigenpose velocity (10 for each). Eigenposes were computed by projecting the flattened 3D marker data for each animal onto the top 10 PCs, defined from performing PCA on the entire dataset of 3D marker data across all sessions. Eigenpose velocity was computed as the temporal difference of the eigenpose. Splines were computed through cardinal spline interpolation following previous procedures^110,111^. In brief, at each time point *t*, the fractional distance α(t) was computed for each feature value to the nearest binned values (which were pre-specified). We set the interpolation parameter s = 0.7. This feature set accommodated nonlinear relationships between neural activity and kinematics, thus providing a highly expressive model with relatively few parameters.

To learn the parameters *W_i_* and *b*_0_, we used MATLAB’s fminunc function to maximize the Poisson log-likelihood of the observed spike train (*n*) given the model spike number 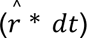 and under the prior knowledge that the parameters should be small through an L2 penalty (with hyperparameter β = 1). Model performance for each cell was computed using the pseudo-R^2^ value, which is the ratio of the increase in log-likelihood of the model compared to the log-likelihood of a mean firing rate model, compared to the increase in log-likelihood of the model compared to the log-likelihood of a fully-saturated model (where there is a parameter for every time point). Performance was quantified through 5-fold cross-validation, where each fold was 20% of the data and chosen from 30-second sections of the entire dataset such that no two folds overlapped. Cells were designated as encoding behavioral kinematics if the pseudo-R^2^ was positive across all folds.

#### Identification of species-typical and task templates

Templates were identified through two methods: one based on the eigenposes, and another based on the accelerometer.

For the eigenpose-based templates, we employed the same eigenposes utilized in fitting the GLM, focusing on the top 5 eigenposes. We identified 1-second long segments within each active (non-idle) behavior, denoted as “snippets.” We then filtered these snippets based on their average variance, discarding those with low variance. Following a methodology similar to how we identified templates for the lesion data, we cross-correlated each snippet with the eigenpose signals throughout the entire recording sessions to identify matches. Candidate matches were those exhibiting a mean correlation coefficient > 0.8 (across all 5 eigenposes) and a mean Euclidean distance < 120 (across all 5 eigenposes). To prevent overlap among matches for a single template, we greedily selected matches, as described above. Snippets were recorded as a template if there were more than 10 matches in the session.

For the accelerometer, the three axes of the accelerometer signal for each recording session are downsampled to 50 Hz and individually z-scored across time. Similar to the procedure described above, we identified 1-second long snippets within each active (non-idle) behavior, filtered based on the average variance to discard those with low variance, and then cross-correlated each snippet with the accelerometer signal from the entire session. Candidate matches were those exhibiting a mean correlation coefficient > 0.75 (across all 3 accelerometer axes) and a mean Euclidean distance < 20 (across all 3 axes).

To combine templates of both types, we took templates that had at least a > 0 correlation across all matches for both accelerometer and eigenpose data. We then removed overlapping templates as above. Templates and their matches were then assigned a ‘task’ label if at least 90% of the matches were within 10 seconds of a lever press, and assigned a species-typical label if 90% of the matches were not within 10 seconds of a lever press.

#### Quantification of spiking activity during template matches

For each template, we considered cells that were active for at least 5 trials and had an average firing rate > 0.1 Hz. For each cell-template pair, we calculated the average correlation in activity across trials by smoothing the single-trial firing rate using a Gaussian filter with a width of 10 bins, and then computing the correlation between every pair of trials. We then performed similar calculations for the average correlation across accelerometer signals and eigenpose signals within each pair of trials. Utilizing this dataset, we computed the slope of both the neural correlation and the accelerometer (or eigenpose) correlation.

To determine whether there was a difference in the task versus species-typical templates in the neural trial-by-trial correlation or the slope values, we fit parallel lines to the data as a function of the accelerometer or eigenpose correlation values and used an F-test determine if the intercept was significantly different.

#### Comparison of GLM-based tuning curves

To compare the tuning curves learned by the GLM, we constructed ‘model-derived tuning curves’ by using the parameters learned in the GLM. These tuning curves return an output similar to standard firing rate based tuning curves, with the exception that the average firing rate for a given variable value has the contribution of other variables regressed out. To compute the tuning curve (*C*) for a variable *i*, we first binned values taken by variable *i* into *M* bins (indexed by *m*; default *M* = 200). We then computed the expected firing rate for each bin using the learned parameters *W_i_*, a feature vector 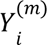 that returns the feature vector assuming that variable *i* takes the value within bin *m*, and scaling variable γ that denotes the average firing rate contribution from all other variables that are significantly encoded by that cell: 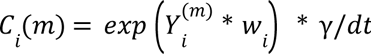. The value of γ is the baseline firing rate multiplied by the average firing rate contribution from every other variable.

To compare the GLM-based tuning curves within species-typical behaviors, we split them into two groups: (1) face groom, anogenital groom, mixed groom, mixed scratch 1, mixed scratch 2, body groom, right scratch, left scratch, fast groom, and (2) high rear slow, high rear fast, fast movement, walk, mid-rear fast, mid-rear look down, mid-rear slow, mid-rear pause, and small movement. The first group consisted entirely of grooms and scratches, which were infrequently used during the task, while the second group consisted of behaviors similar to those more often used in the last, like rearing and fast movements. This split also generated groups similar in the number of time points.

#### Construction of postural tuning curves

For each cell that was active and well-described by the GLM (and thus a function of the behavioral kinematics) in either domain, we first identified the eigenpose value (for the top 5 eigenposes) for which the cell was most active. To do this, we binned the data into 12 Hz bins, sorted the bins according to firing rate, took the top 100 bins, clustered the eigenpose values based on Euclidean distance (using the clusterdata function in MATLAB), and identified the ‘preferred eigenpose’ as the most common pose. We then computed the Euclidean distance between the eigenpose at each timepoint and the preferred eigenpose, and then computed the firing rate as a function of this distance. This procedure was performed initially within a domain only (i.e., with the task or species-typical domain); activity was then computed as a function of eigenpose distance for all time points in the opposite domain.

This procedure was also performed for two consecutive time points, i.e. the ‘preferred eigenpose’ was a consecutive series of two eigenposes.

#### Statistical tests

Unless otherwise noted, statistical tests were nonparametric and two-sided. Multiple comparison tests were used where justified. In figures and text, p-values less than 0.0001 were written as p < 0.0001.

## Acknowledgements

We thank members of the Ölveczky Lab for discussions and comments on the manuscript. We also thank Diego Aldarondo for help with DANNCE-related data and processing. This work was supported by NIH grants 5R01NS099323-07 toB.P.Ö., by a Jane Coffin Childs Memorial Fund Fellowship and Harvard Brain Science Postdoc Pioneer Award to K.H., and by a Helen Hay Whitney Fellowship and a NIH K99 to J.D.M.

