## Supplemental Figures for "Differential kinematic coding in sensorimotor striatum across species-typical and learned behaviors reflects a difference in control"

**Supplemental Figure 1**

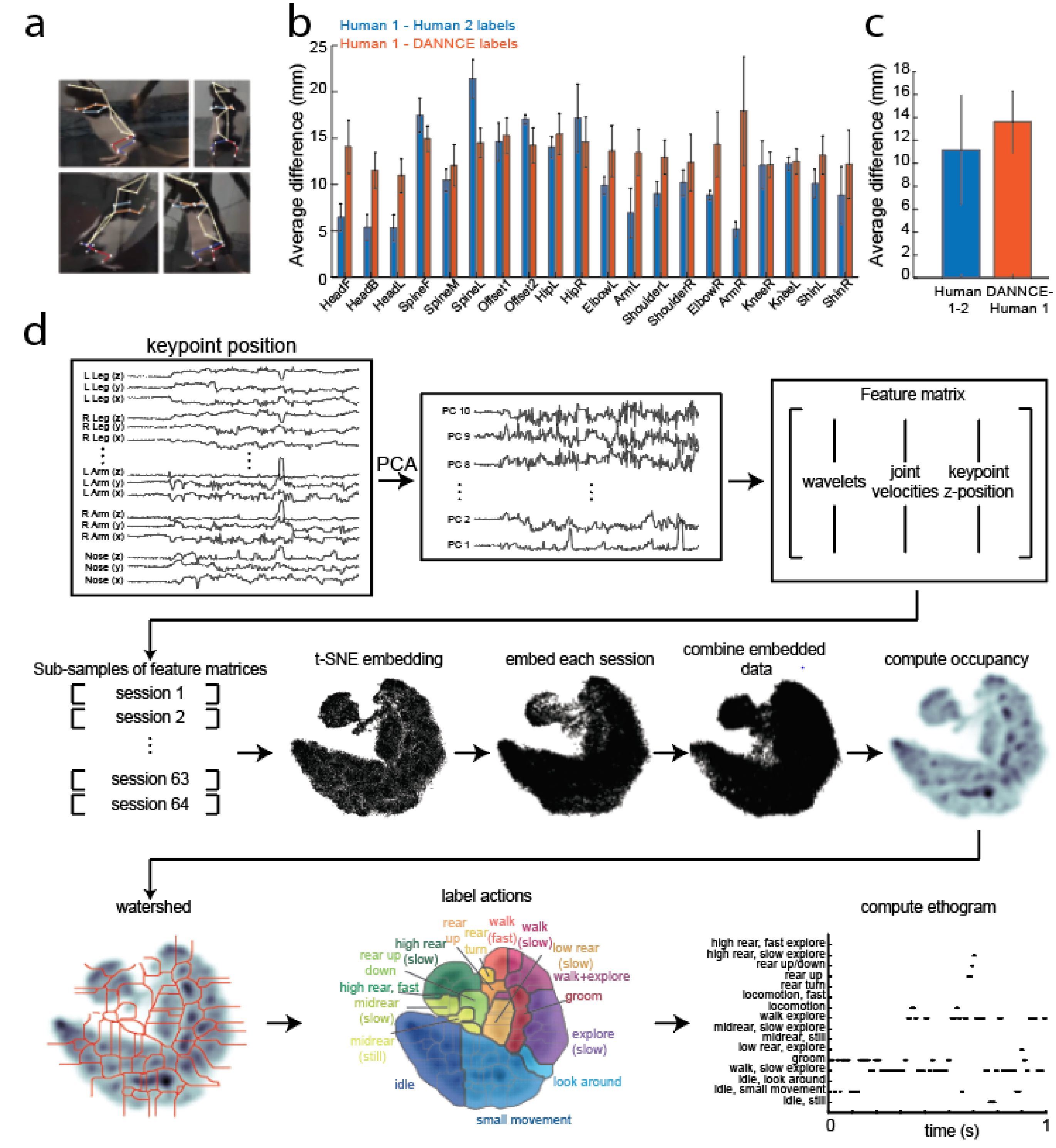


**Supplemental Figure 1. Animal pose tracking with DANNCE and characterization of animal behavior using MotionMapper. a.** Viewpoints from 4 of the 5 cameras for a single point in time. The colored skeleton superimposed on the animal represents the DANNCE keypoints. **b.** The average difference in millimeters (mm) between the 3D positions of DANNCE keypoints (in orange) and those obtained from two expert human labelers (in blue). A total of 80 frames were labeled for comparison; these labels were not used to train the DANNCE network. Error bars represent standard deviation. **c.** Same data as in B, but averaged across keypoints. Error bars represent standard deviation. **d.** A schematic of the MotionMapper pipeline adapted for 3D keypoint data (Marshall et al. 2021). First we use DANNCE to track keypoints for each session. PCA is then applied to all keypoints from all sessions to obtain the top 10 principal component axes; all data is then projected onto these axes. We then construct a feature matrix composed of wavelet transforms of the PC-projected data, velocity of each keypoint, and z-position (height) of each keypoint. We then define a low-dimensional space (the 'behavioral map') upon which the rows of the feature matrix can be embedded. To do this, we first selectively subsample from the feature matrix across sessions and then embed these points into a 2-dimensional space using t-SNE. The entire feature matrix is subsequently re-embedded into this low-dimensional space. To cluster regions of this space into discrete 'finely-parsed' behaviors, we smooth to get a density of points across the space and use watershed clustering to segment regions. We then label each segment manually, and group the finely-parsed regions into a smaller set of categorically-distinct 'coarsely-parsed' behaviors. Each coarsely-parsed behavior is denoted by a distinct color. We can then use this map to define an ethogram, which is the behavior expressed by the animal at every point in time.

### Supplemental Figure 2

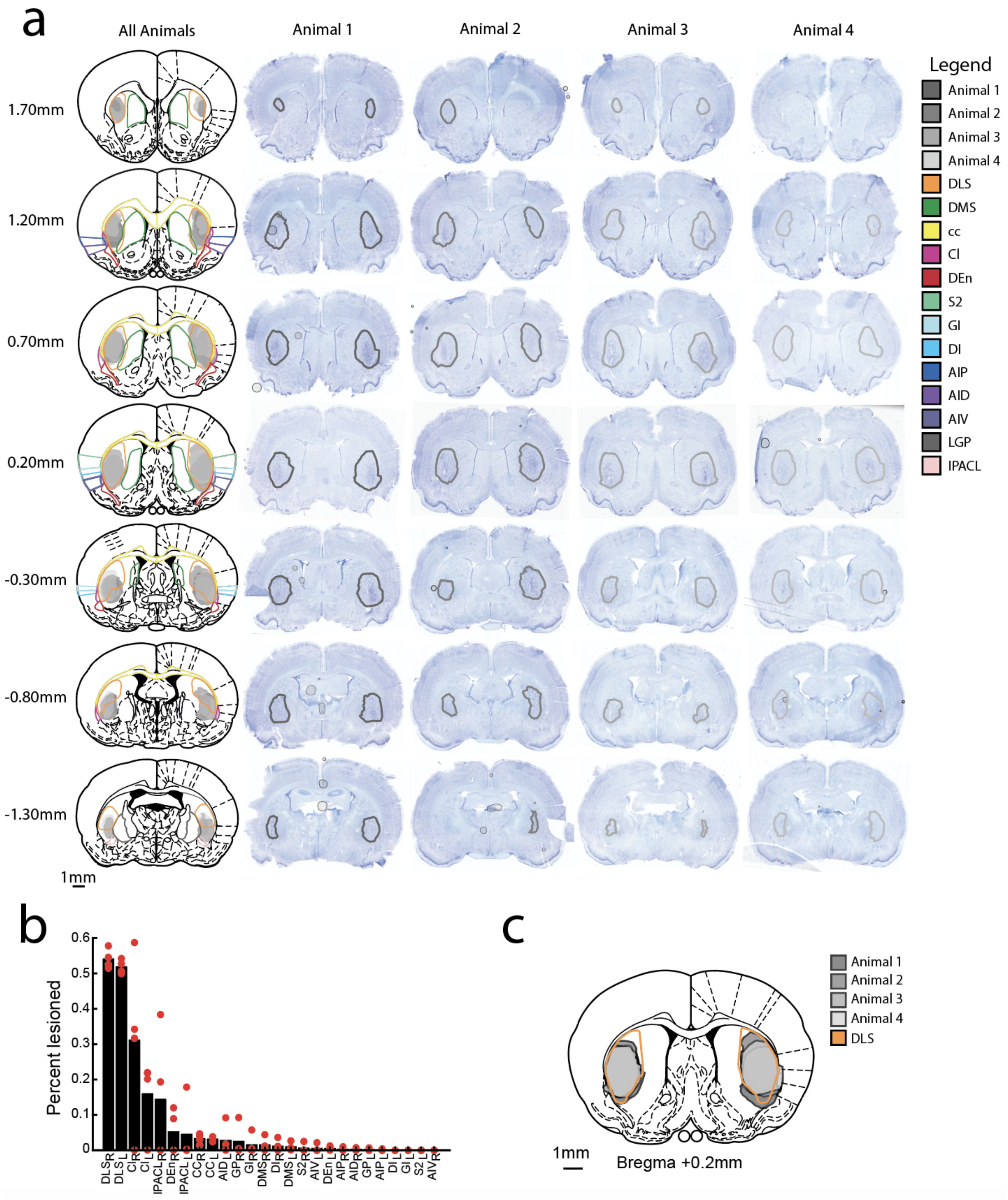

**Supplemental Figure 2. Quantification of the bilateral DLS lesions. a.** The leftmost column displays histological images illustrating the lesion boundaries (shaded regions) for four animals at different distances from Bregma. The distinct regions are outlined and color-coded following the legend on the right. The abbreviations for each region are as follows (Paxinos and Watson 1998): dorsolateral striatum (DLS), dorsomedial striatum (DMS), corpus callosum (cc), claustrum (Cl), DEn (dorsal endopirifom cortex), secondary somatosensory cortex (S2), granular insular cortex (GI), dysgranular insular cortex (DI), agranular insular cortex posterior (AIP), agranular insular cortex dorsal (AID), agranular insular cortex ventral (AIV), lateral globus pallidus (LGP), interstitial nuclei of posterior limb of ac, la (IPACL). DMS and DLS boundaries are derived from (Dhawale et al. 2021). The fourmost right columns depict the lesion boundaries for each animal. **b.** The graph displays the fraction of each region that has been lesioned for each animal. Each red dot corresponds to one animal, and the black bars represent the average fraction across all animals. Note that substations portions of DLS on both sides are lesioned. **c**. Composite image showing the DLS lesion boundary for all animals (+0.2 anterior to Bregma).

### Supplemental Figure 3

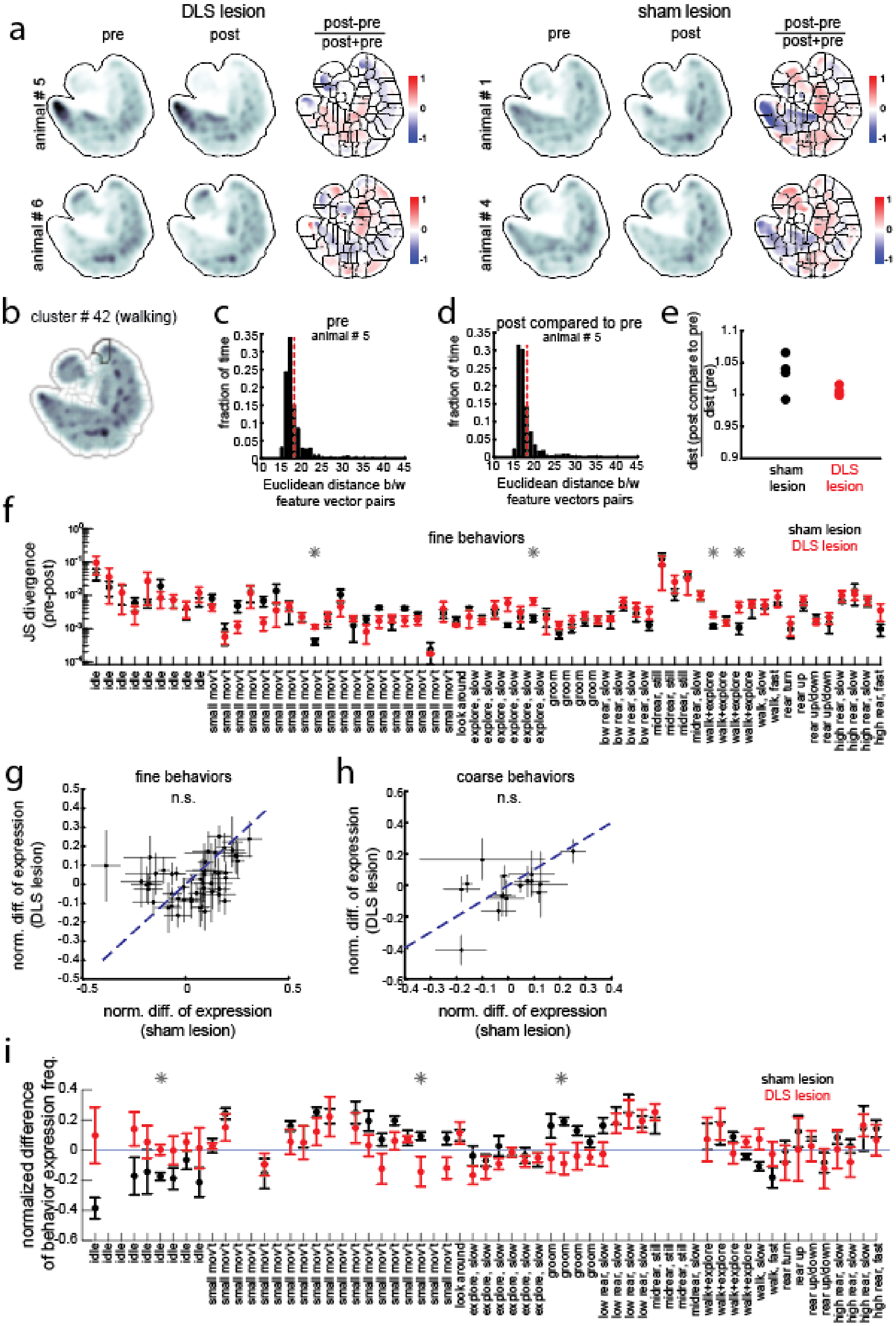

**Supplemental Figure 3. Comparison of behavioral kinematics before and after DLS lesions. a.** Two example animals' behavioral maps before and after DLS lesions (left) or sham lesions (right). These maps are generated by averaging all four sessions before or after surgery. The colored maps indicate the normalized differences across maps, highlighting alterations in behavioral patterns following surgery (and recovery time) in both cohorts. **b.** The behavioral map, averaged across all sessions for both cohorts, with the fine clustering overlaid. A single cluster (#42) is outlined, and serves as the focus of the analysis in panel C-E. **c.** The Euclidean distance between pairs of feature vectors (see Methods) for a specific animal prior to surgery. Feature vectors were averaged across a maximum of 10 consecutive samples during a single 'bout' into the behavioral cluster to reduce the number of pair comparisons. **d.** Similar to C, but the Euclidean distance is computed between feature vectors after surgery and feature vectors before surgery. **e.** This plot shows the normalized Euclidean distance between feature vectors (panel D / panel C) for all animal in both cohorts, for cluster #42. The data presented in Fig 2D is the average across animals in each cohort, for all behaviors. **f.** The Jensen-Shannon (JS) divergence of embedded features within each fine behavioral cluster is shown for DLS-lesioned animals (red) and sham-lesioned animals (black). Error bars represent SEM. Black stars indicate the behaviors where the JS divergence of DLS-lesioned animals exceeded that of the sham-lesioned animals. However, note that only 6% (4/62) show significant differences at $p < 0.05$ (uncorrected), which is approximately chance level. **g.** The normalized difference in expression across finely parsed behaviors before and surgery for sham-lesioned animals and DLS-lesioned animals (same data as in I). Errorbars correspond to SEM across animals. There was not a significant difference in the magnitude (absolute value) of change between cohorts (signrank $p = 0.095$). **h.** Same as G, but for coarsely parsed behaviors (data same as Fig 2H). There was not a significant difference in the magnitude of change between cohorts (signrank $p = 0.35$). **i.** The normalized change in behavioral expression before and after surgery, quantified as (post-pre)/(post+pre). Error bars represent the SEM across animals. Finely-parsed behaviors with few examples ( < 5 seconds) were ignored for this analysis. In total, 3/53 (5%) behaviors were significantly different across sham and DLS-lesioned animals, which is chance level for $p = 0.05$.

**Supplemental Figure 4**

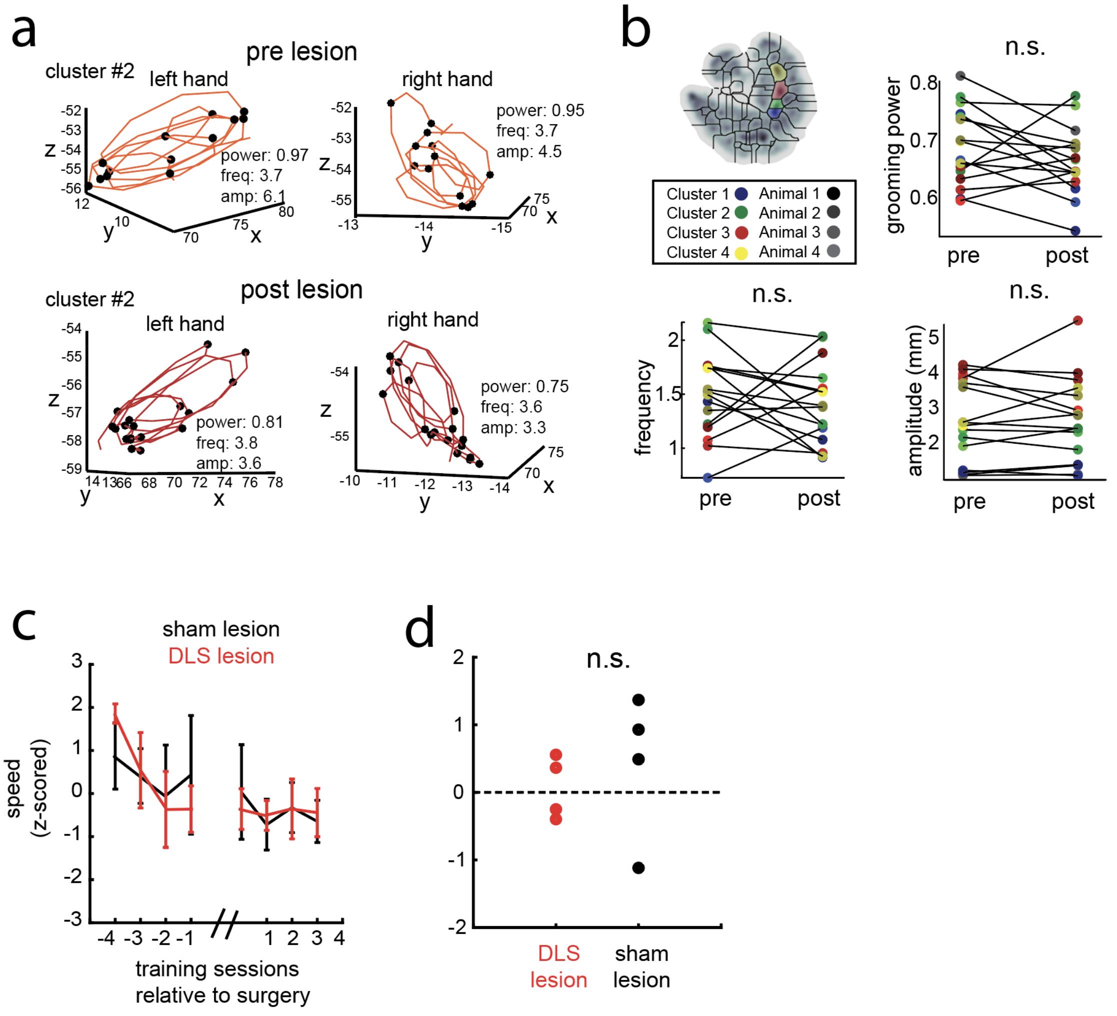


**Supplemental Figure 4. Lesioning DLS does not induce changes in vigor. a.** An example grooming bout, showing the right and left wrist positions for an animal before and after the lesion. Grooming bouts were identified as 1-second bouts in one of the 4 grooming clusters identified through motion mapper. Numbers on the right side of each panel correspond to grooming power (the power in the 2-6 Hz band divided by the 0-14 Hz band), grooming frequency (detected via peak/troughs, denoted by black stars), and grooming amplitude (also computed via peak/troughs). **b.** Top-left: the four analyzed grooming clusters along with the legend for different grooming clusters and animals. Top-right: The difference in grooming power across grooming clusters and animals. There was no significant difference after DLS lesion using a linear mixed effects model ($p = 0.88$, see Methods). Bottom-left, same at the top-right, but for the frequency (linear mixed effect model, $p = 0.86$). Bottom-right, same as the top-right, but for the amplitude (linear mixed effects model, $p = 0.99$). **c.** The speed, z-scored across animals. Errorbars correspond to SEM. **d.** There was no significant difference across cohorts in the difference of speed from the last day prior compared to the first day after surgery ($n = 4$ in each cohort, ranksum $p = 0.49$).

### Supplemental Figure 5

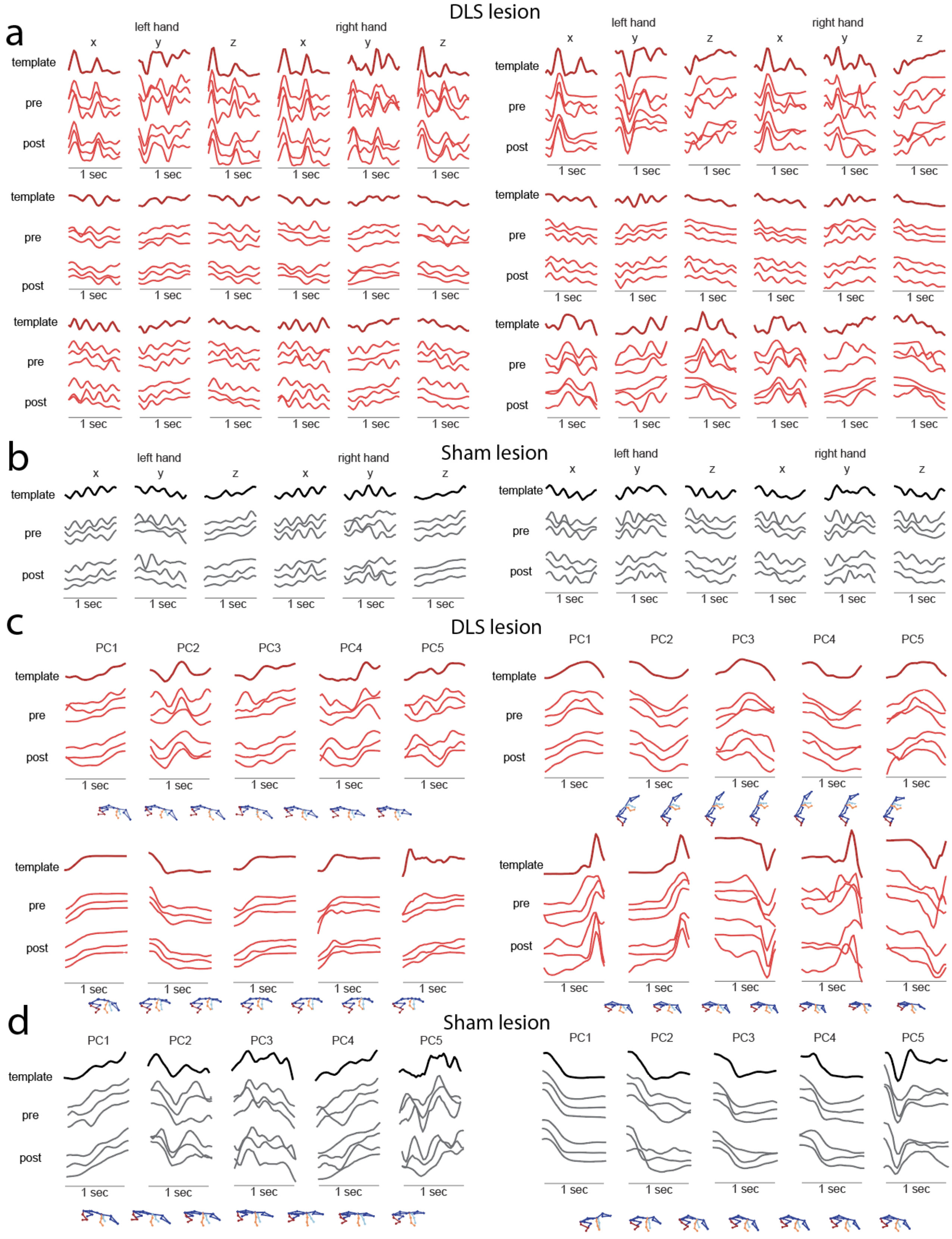

**Supplemental Figure 5. Example grooming and whole-body templates, and their matches, before and after DLS or sham lesions. a.** Six example grooming templates and three example matches per template. Templates are shown in dark red, with pre-lesion and post-lesion matches in lighter red. The x, y, and z positions of both hands, which are used to identify templates and matches, are shown for each example. **b.** Same as A, but for sham-lesioned animals. Templates are in black with matches in gray. **c.** Same as A, but for whole-body templates. Signals shown are the first five eigenposes (denoted by PCs). The skeletons on the bottom denote the whole-body movement captured by the template. **d.** Similar to C, showing the first five eigenposes of whole-body templates and their corresponding matches for sham-lesioned animals.

**Supplemental Figure 6**

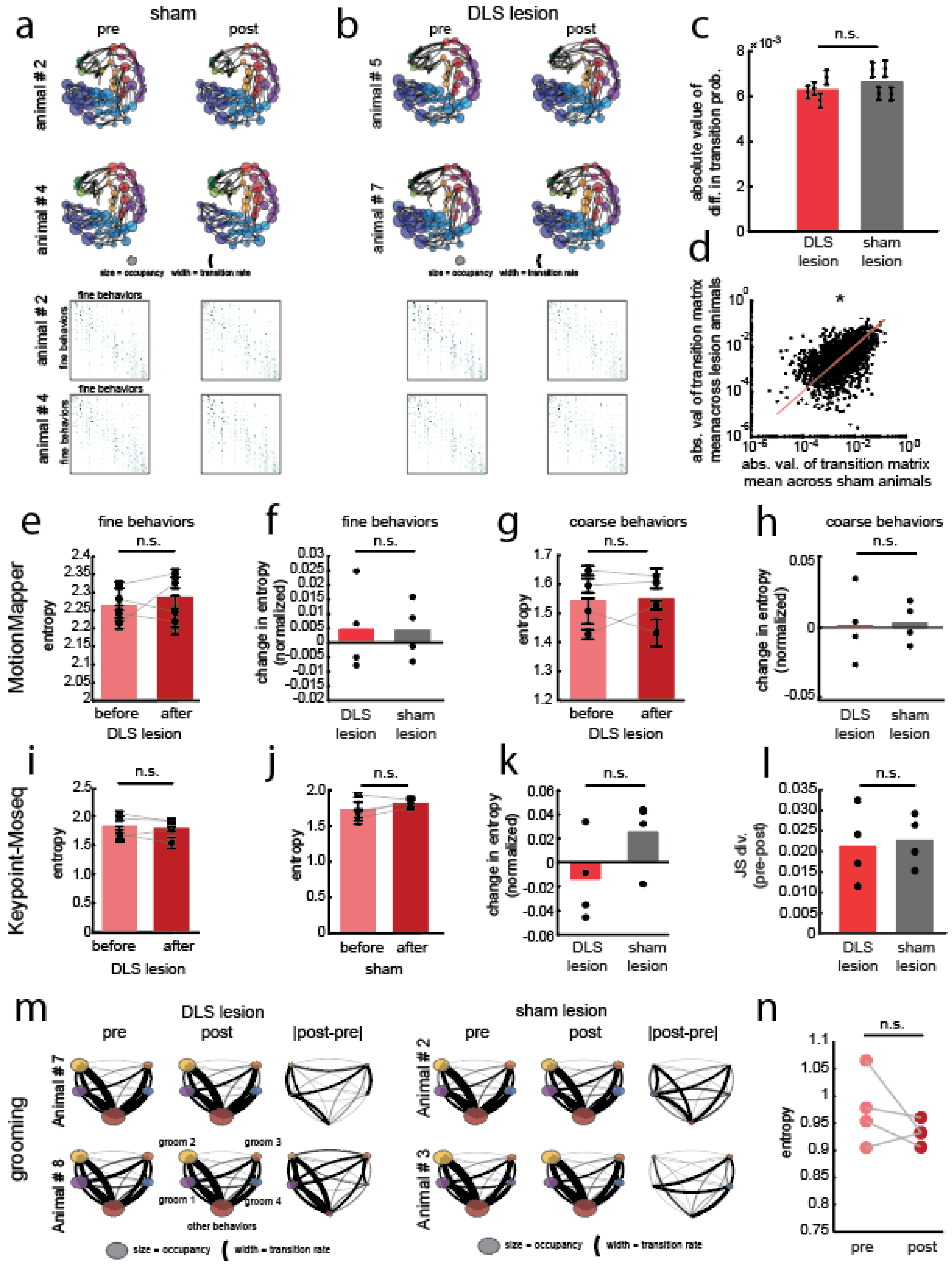

**Supplemental Figure 6. Lesioning DLS does not change the transition structure of behavioral sequences. a.** Top two rows: occupancy and transition of two example sham lesioned animals before and after surgery. Each circle corresponds to a finely parsed behavior, with the size of the circle corresponding to the occupancy of that behavior. The width of each line between behaviors denotes the transition from one behavior to another. The color corresponds to the coarse label. Bottom two rows: the transition matrix for the same data as in the top two rows. Darker colors correspond to more commonly-taken transitions. **b.** Same as A, but for DLS-lesioned animals. **c.** There was no significant difference between cohorts in the average magnitude of change in the transition matrix (ranksum p = 0.69, n = 4 in each cohort). Errorbars correspond to the mean and SEM for each animal across all transition matrix values. **d.** The absolute value of the difference between transition matrices, averaged across animals in each cohort. Each point corresponds to one of the $62^2$ transitions. The median absolute difference before/after surgery was larger for sham-lesioned animals compared to DLS-lesioned animals (signrank p = 0.01), meaning that sham-lesioned animals exhibited more differences after surgery than DLS-lesioned animals. **e.** The entropy of the transition matrix for finely-parsed behaviors before and after DLS lesions, computed as in (Markowitz et al. 2018; Wiltschko et al. 2015)**.** Each point corresponds to one animal, with error bars denoting the SEM across the four sessions. There is no significant difference before/after DLS is lesioned (n = 4, signrank p = 0.63). **f.** Comparison of the modulation [(post-pre)/(post+pre)] in entropy for DLS-lesioned and sham-lesioned animals. There is no significant difference across groups (n = 4 in each group, ranksum p = 0.89). **g.** Same as D, but for coarse behaviors. There was no difference in entropy after DLS was lesioned (signrank p = 0.88). **h.** Same as E, but for coarse behaviors. There was no difference across groups (signrank p = 0.89). **i.** The entropy across syllables before and after DLS was lesioned, as detected via Keypoint-Moseq (Weinreb et al. 2023) (see Methods for details). There were 16-33 frequently-used syllables across all animals. Similar to D and F, there was no difference in the entropy after DLS was lesioned (signrank p = 0.38). **j.** Same as H, but for sham-lesioned animals. There was no difference after the sham lesion (signrank p = 0.25). **k.** Same as E or G, but for the modulation in entropy computed using Keypoint-Moseq syllables detected in DLS- or sham-lesioned animals. There was no difference across cohorts (ranksum p = 0.2). **l.** The JS divergence in the distribution of syllables in DLS-lesioned or sham-lesioned animals before and after lesion. There was no difference in the JS divergence across cohorts (ranksum p = 0.89), consistent with our prior results using MotionMapper. **m.** The average transition probability between the four grooming clusters. The fifth node (on the bottom) corresponds to all other behavioral clusters. Larger node size denotes high occupancy, and thicker transition lines denote increased transition rate (ignoring self-transitions). Note that while there are changes in the transition rate between grooming behaviors for DLS-lesioned animals, there are changes of similar magnitude for sham-lesioned animals. **n.** The entropy in the grooming-specific transition matrix before and after DLS lesion. There was no difference after lesion (n = 4, signrank p = 0.38).

**Supplemental Figure 7**

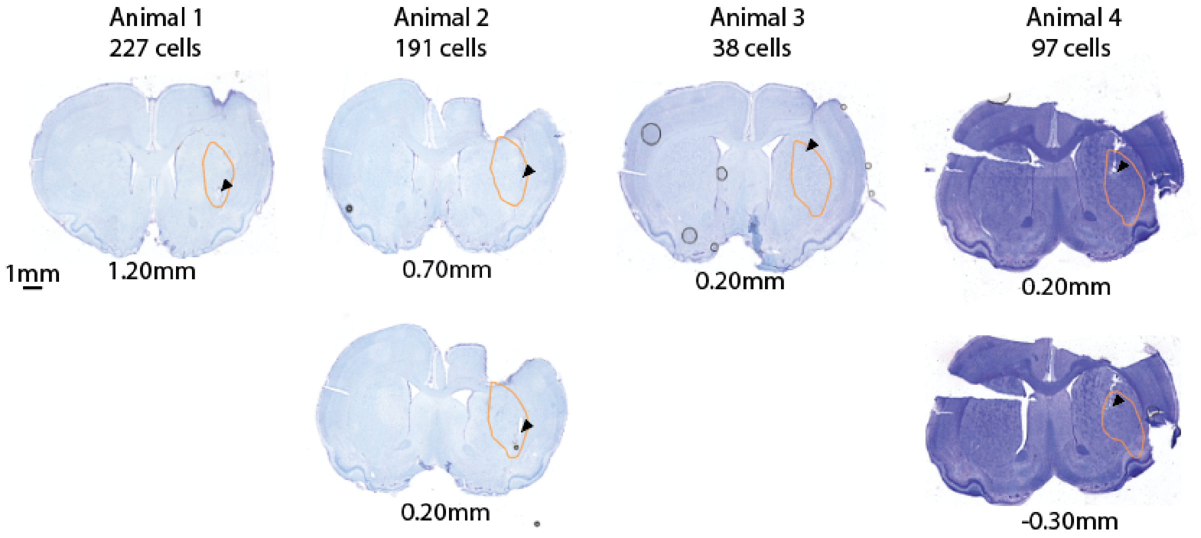


**Supplemental Figure 7. Histology for tetrode implantation in four animals.** The number of cells in the presented dataset from each animal are listed beneath each animal number. The orange lines indicate the DLS boundary, as given in (Dhawale et al. 2021). The black arrow indicates the location of the endpoint of the tetrode bundle. The number beneath each image corresponds to the distance from Bregma.

**Supplemental Figure 8**

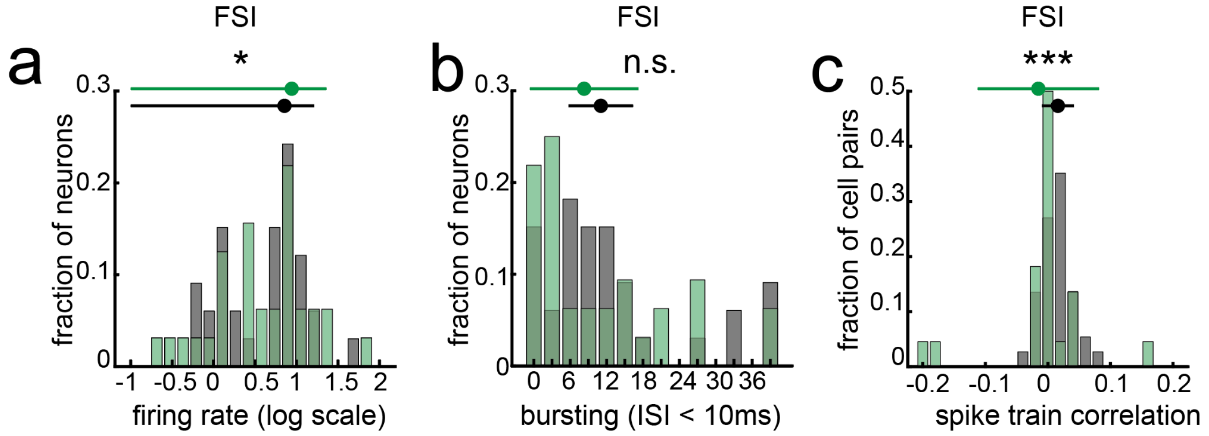


**Supplemental Figure 8. FSIs exhibit a wider range of firing rates and correlated spike trains, but not a wider range of bursting, for exploratory species-typical versus task behaviors. a.** The distribution of mean firing rates for neurons active in either domain (in a shaded portion of panel d) during the species-typical behavior (gray) or the task (green). Plotted above the distribution are the mean +/- standard deviation of the distribution. The distribution of firing rates during the task had a significantly larger variance than during species-typical behavior (n = 33 for both groups, F-test p = 0.015). **b.** The distribution of bursting, quantified as the percentage of interspike intervals less than 10 ms, for activity during the species-typical behavior (gray) or the task (green), with the mean +/- standard deviation plotted above as in d. The variance during the task was not significantly larger than during species-typical behavior (n = 33 for both groups, F-test p = 311). **c.** The distribution of correlation coefficients computed between smoothed spike trains across all cell pairs, with the mean +/- standard deviation plotted above. For each domain, only cells active in that domain were considered. The distribution of correlation coefficients during the task had a significantly larger variance than during species-typical behavior (task n = 37, species-typical n = 27, F-test p < 0.0001).

**Supplemental Figure 9**

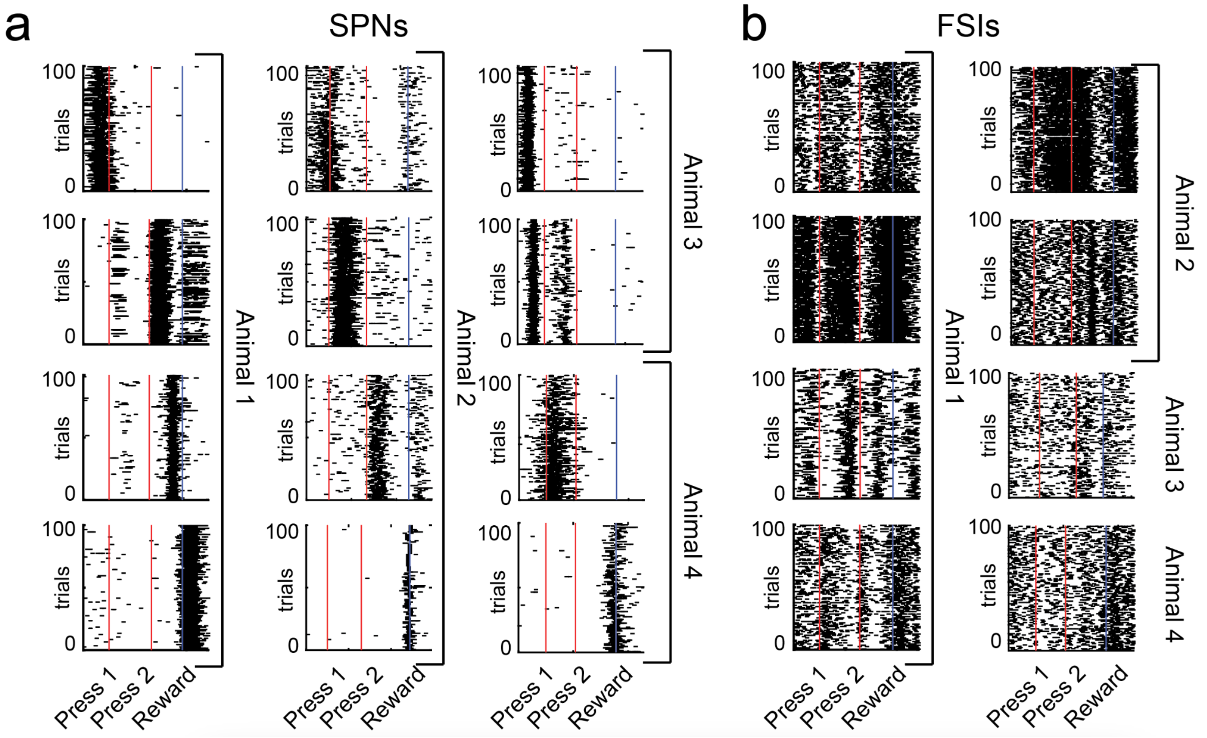


**Supplemental Figure 9. Spiking patterns during the timed lever-pressing task. a.** Spiking activity from representative SPNs of each animal while they executed the timed lever-pressing task. Red lines correspond to each tap, while the blue line indicates reward. Trials were time-warped. Each cell showed stable activity across trials (the firing pattern across trials was more correlated than the top 5% of a null distribution created by circularly shifting the spike trains in time). The 100 trials shown represent those most closely aligning with that cell's average firing pattern during the task. Note that the firing patterns form a sequence, with most cells firing during specific times of the task. **b.** The same as A, but for FSIs. Note that these cells exhibit less sparse (dense) firing patterns, as they tend to be active throughout the entire task period.

### Supplemental Figure 10

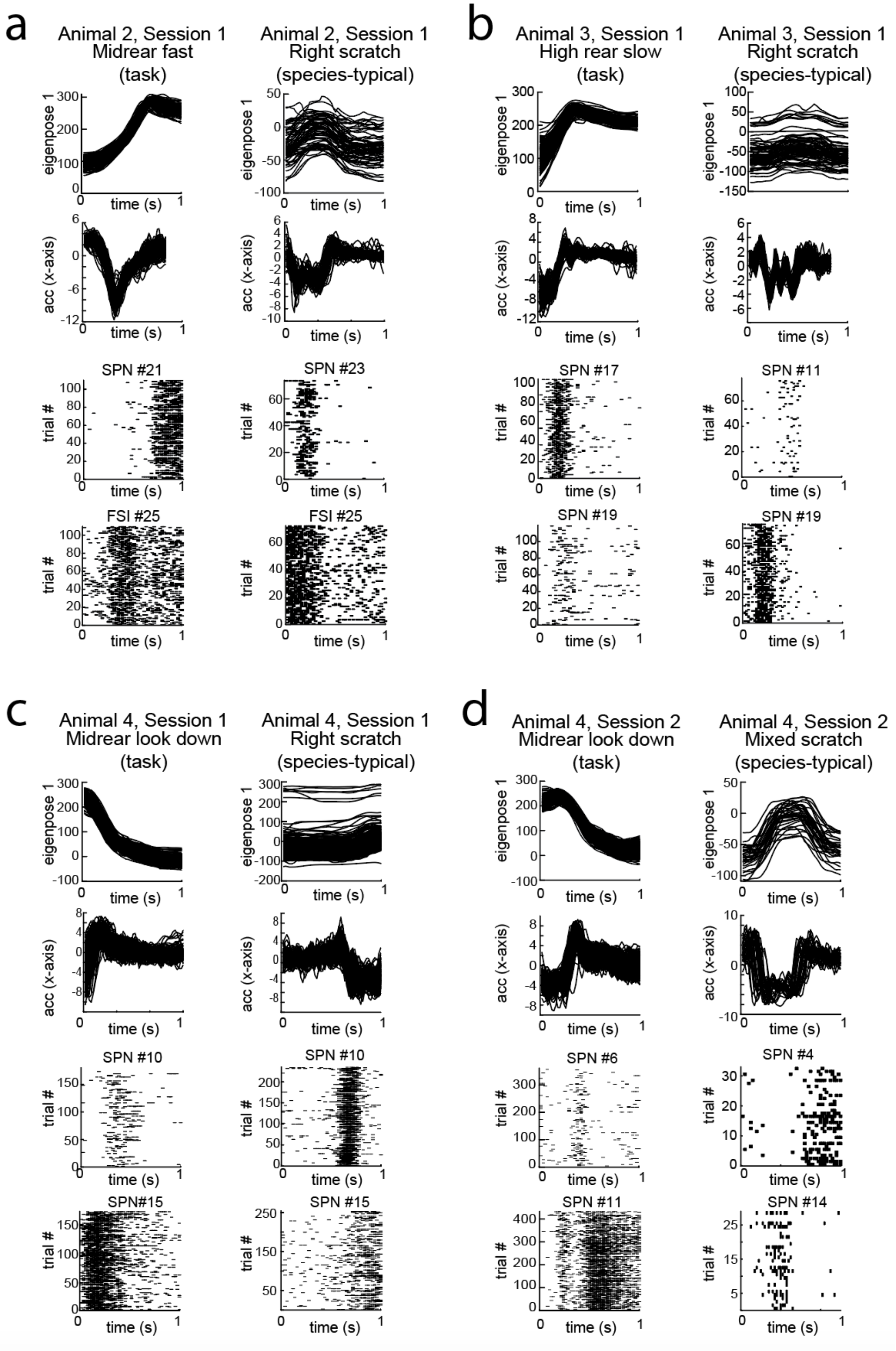

**Supplemental Figure 10. Comparison of neural activity during movement templates identified during the task or during spontaneous species-typical behavior. a.-d.** Neural activity from SPNs and FSIs during species-typical and task-derived templates for three different animals and sessions. The top two rows of each plot correspond to the eigenpose and accelerometer of the template matches. The bottom two rows correspond to the raster plots for each cell during the same time the matches are expressed. Cell numbers (which are specific to each session) are given above each relevant subplot. Note that in some cases, the same cells are plotted for task and species-typical templates.

### Supplemental Figure 11

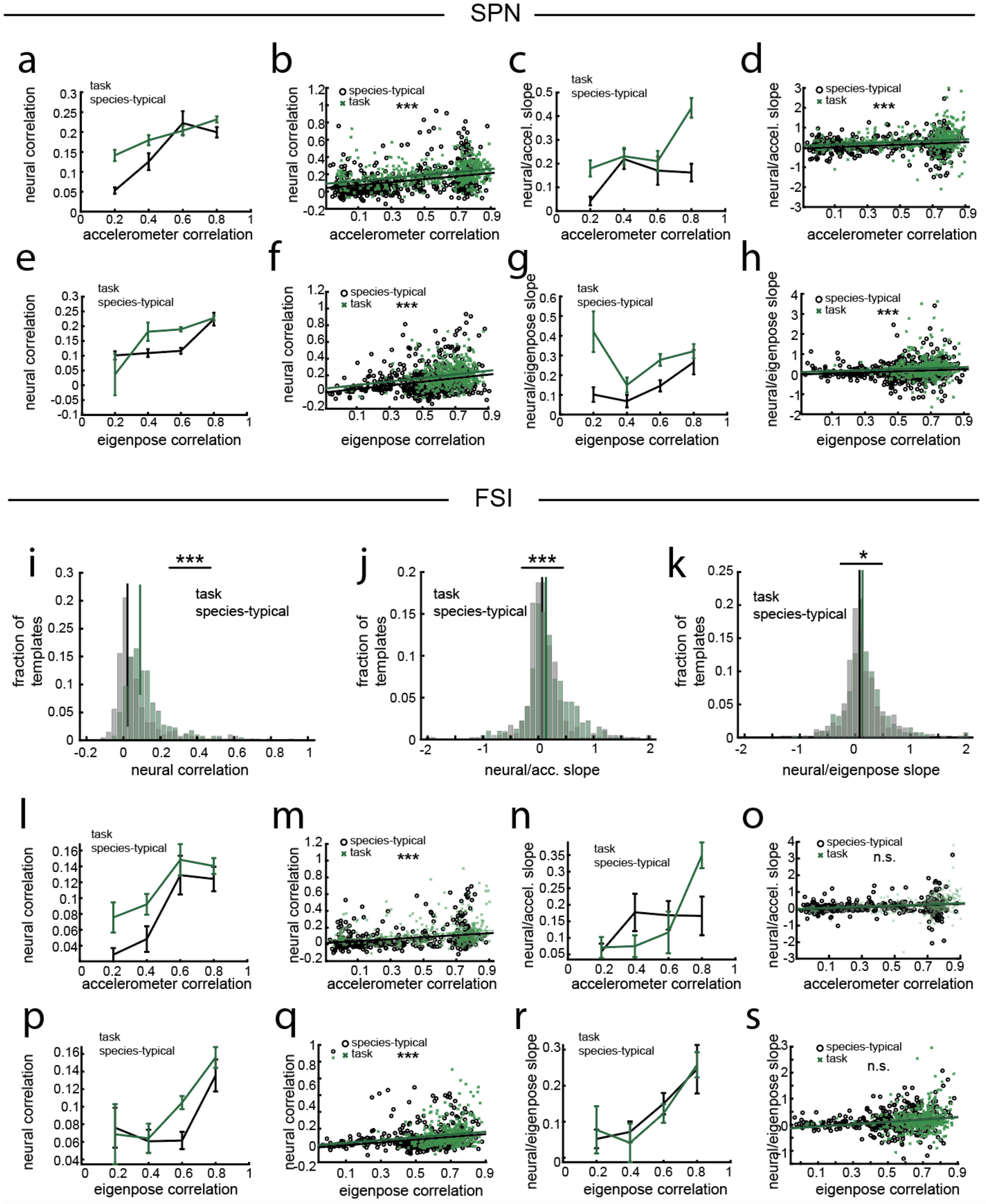

**Supplemental Figure 11. Comparison of SPN and FSI activity during templates identified during the task or during spontaneous species-typical behavior. a.** The average across-trial neural correlation plotted against the across-trial accelerometer correlation, averaged across SPNs for each task-specific template and species-typical template. **b.** The raw data that generated B, with lines of parallel slope fit to each group. There was a significant difference in the intercept ($p = 3*10^{-8}$). **c.** The average across-trial neural correlation plotted against the slope between the across-trial accelerometer correlation and across-trial neural correlation, averaged across SPNs for each task-specific template and species-typical template. **d.** These plots show the raw data that generated C, with the best-fit, parallel lines overlaid. There was a significant difference in the intercept ($p = 2*10^{-7}$). **e.** Similar to A, but for the correlation in the eigenposes rather than the accelerometer. **f.** Similar to B, but for the correlation in the eigenposes rather than the accelerometer. There was a significant difference in the intercept ($p = 2*10^{-6}$). **g.** Same as C, but for eigenpose instead of accelerometer. **h.** Same as D, but for eigenpose instead of accelerometer. There was a significant difference in the intercept (p =0.0003). **i.** The distribution of the average correlation in FSI activity across pairs of trials for task-specific templates (green) and species-typical templates (black). The average across-trial neural correlation was lower for species-typical templates compared to task templates (task-specific template n = 122, species-typical template n = 278, ranksum $p < 0.001$). **j.** The distribution of the slope between the average across-trial FSI activity correlation for each task-specific or species-typical template, where the across-trial FSI activity correlation is the average correlation in the firing rate across all pairs of trials, and the average across-trial accelerometer correlation, which is the mean correlation across accelerometer signals for all trial pairs. When averaging across neurons for each template, we observe that the slope is much higher for task-specific templates (ranksum $p < 0.00004$). **k.** The same as panel J, but for eigenpose signals instead of accelerometer signals. Again, we observe that the slope is much higher for task-specific templates (ranksum $p < 0.018$). **l.** The average across-trial neural correlation plotted against the across-trial accelerometer correlation, averaged across FSIs for each task-specific template and species-typical template. **m.** The raw data that generated L, with lines of parallel slope fit to each group. There was a significant difference in the intercept (p = 0.0009). **n.** The average across-trial neural correlation plotted against the slope between the across-trial accelerometer correlation and across-trial neural correlation, averaged across FSIs for each task-specific template and species-typical template. **o.** These plots show the raw data that generated N, with the best-fit, parallel lines overlaid. There was not a significant difference in the intercept (p = 0.25). **p.** Similar to L, but for the correlation in the eigenposes rather than the accelerometer. **q.** Similar to M, but for the correlation in the eigenposes rather than the accelerometer. There was a significant difference in the intercept (p = 0.006). **r.** Same as N, but for eigenpose instead of accelerometer. **s.** Same as D, but for eigenpose instead of accelerometer. There was not a significant difference in the intercept (p = 0.52).

### Supplemental Figure 12

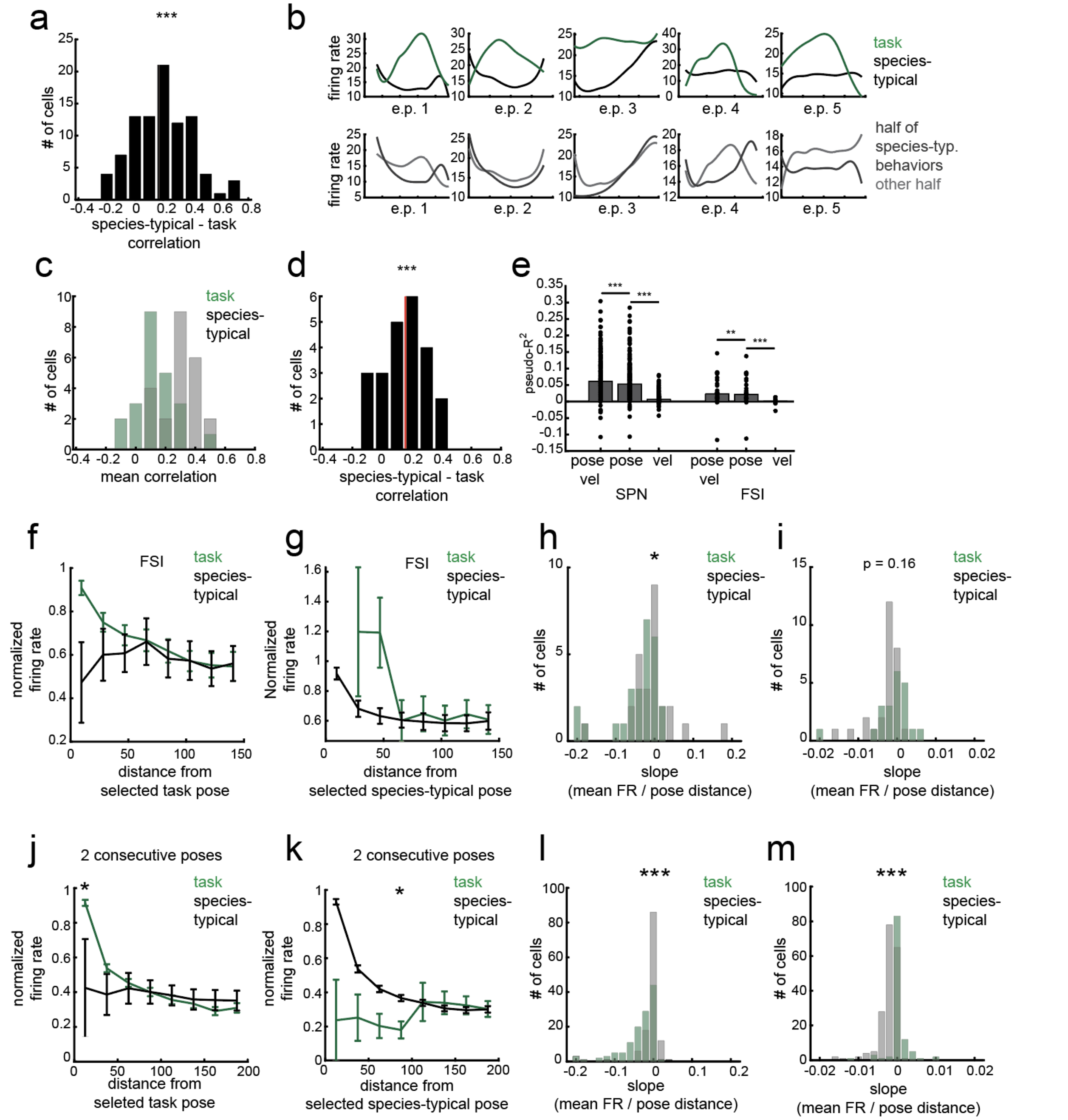


**Supplemental Figure 12. Comparison of kinematic representations during the task with that during spontaneous species-typical behavior. a.** Histogram of the difference in the mean model-derived tuning curve correlation coefficient between halves of species-typical behaviors and the task and species-typical behavioral domains (i.e., between the green and black points forming the histogram in Figure 6C). The red line corresponds to the median of the distribution; it is shifted to the right of 0 with signrank $p < 0.0001$, $n = 89$). **b.** Example model-derived tuning curves for an FSI that significantly encoded pose and pose velocity during the task and species-typical behavior. Top row: model-derived tuning curves computed from task or species-typical behaviors. Bottom row: model-derived tuning curves computed across two evenly-split halves of the species-typical behaviors (similar to Figure 6B). **c.** Histogram of the average correlation coefficient between model-derived tuning curves in each behavioral domain (green), or in evenly divided halves of species-typical behaviors (black) for FSIs (n = 29). **c.** Histogram of the difference in the mean model-derived tuning curve correlation coefficient between halves of species-typical behaviors and the task and species-typical behavioral domains (i.e., between the green and black points forming the histogram in panel C). **e.** Prior results have indicated that the posture of the animal is more predictive of neural spiking than the velocity during the timed-lever pressing tas (Dhawale et al. 2021).To see if this was true for species-typical behavior, we computed the average pseudo-$R^2$ of the three different GLMS on five-folds of held-out data for active SPNs (left) and FSIs (right) during species-typical behavior. The models differ in their features: one uses both pose and velocity features (as elsewhere in the paper), one uses only pose information, and one uses velocity information. For both SPNs and FSIs, the pose + velocity model out-performs the pose-only model (SPN: n = 213, signrank p < 0.0001, FSI: n = 33, signrank p = 0.002), which outperforms the velocity-only model (SPN: n = 213, signrank p < 0.0001, FSI: n = 28, signrank p < 0.00001). **f.** Similar to Fig 7E, but for FSIs. The average normalized firing rate (divided by the maximum value of the task-derived data, mean +/- SEM) is plotted as a function of the Euclidean distance from the selected pose. This panel shows FSIs that were task-tuned, as determined through the GLM (n = 26). **g.** Similar to panel E (and Fig 7H), but the ‘selected’ poses are computed for behaviors in the species-typical behavioral domain. The cells considered are the FSIs that were species-typical tuned, as determined through the GLM (n = 28). **h.** The average slope of firing rate versus the Euclidean distance from the task-selected pose (i.e., the slopes of the non-normalized behavioral tuning curves in panel E). The slope of the cells during the task domain are comparatively lower than in the species-typical behavioral domain (signed rank test p = 0.042). **i.** Same as panel G, but for the Euclidean distance of species-typical selected poses, and for species-typical active FSIs as determined by the GLM (i.e., the slopes of the non-normalized behavioral tuning curves in panel F). The slope of the cells during the species-typical behavioral domain is lower than in the task behavioral domain, but the effect is not significant (signed rank test p =0.16). **j.** Same as Figure 7E, but considering the Euclidean distance from a sequence of two consecutive poses. **k.** Same as Figure 7H, but considering the Euclidean distance from a sequence of two consecutive poses. **l.** Same as Fig 7F, but considering the Euclidean distance from a sequence of two consecutive poses (i.e., computed from data in panel I). The slope of the cells during the task behavioral domain are comparatively lower than in the species-typical behavioral domain (signed rank test p < 0.0001). **m.** Same as Fig 7I, but considering the Euclidean distance from a sequence of two consecutive poses (i.e., computed from data in panel J). The slope of the cells during species-typical behavioral domain are comparatively lower than in the task behavioral domain (signed rank test p < 0.0001).
